# Enhancer RNA Transcription Near Segmentation Gene Enhancers Can Be Analyzed *In Situ* Using FISH

**DOI:** 10.64898/2026.03.18.712550

**Authors:** Christine Mau, Benjamin Schmid, Ezzat El-Sherif

## Abstract

Enhancer RNAs (eRNAs) are non-coding transcripts produced at enhancer regions, which appear to be involved in transcriptional regulation. Up to date, these have been primarily investigated using labor-and cost-intensive genomic techniques. However, the precise mechanisms by which eRNA transcription or the eRNA transcripts themselves mediate transcriptional regulation remain unclear. Here, we present a novel experimental approach that allows us to analyze the characteristics of eRNA transcription in fixed and live whole *Drosophila melanogaster* embryos. We employ the anterior-posterior patterning genes as a model system to investigate the dynamics of eRNA expression, utilizing an imaging-based approach. We combined high-sensitivity fluorescence *in situ* hybridization (FISH) chain reaction (HCR) with high-resolution confocal microscopy to detect eRNA and mRNA molecules. Through this experimental assay, we identified foci of elevated transcriptional activity that generate eRNA transcripts correlated with mRNA production at the same gene locus. We could show that this eRNA transcription is independent of promoter activity. Additionally, we demonstrate that insulators can influence eRNA transcription, resulting in loss of eRNA transcription. Moreover, we observe that eRNAs can originate both within classical enhancer regions and outside of them, including from foreign bacterial sequences when these are placed near enhancer sequences, underscoring the strong influence of local regulatory context on eRNA initiation. In live embryos using MS2-MCP live imaging, our analysis of insulators showed a modest reduction in mRNA burst intensity accompanied by a slight increase in burst frequency. Overall, our imaging-based approach offers a novel platform for dissecting enhancer–eRNA interactions and could be adapted for wider applications.

## Introduction

In higher eukaryotes, enhancers are regulatory DNA elements that do not directly activate gene expression but instead modulate the activity of promoters, which function as molecular valves to recruit RNA Polymerase II (Pol II) and initiate mRNA synthesis (Ryu et al., 2024; Lenhard et al., 2012). Despite extensive study, a comprehensive model explaining how enhancers control gene activity and determine transcriptional output remains elusive (Ryu et al., 2024; Zhang et al., 2024; Li et al., 2023).

Recent discoveries, however, provide new avenues for understanding enhancer function. First, transcription is now recognized to occur in bursts, with gene promoters alternating between active and inactive states, resulting in episodic mRNA production. Although enhancers appear to influence transcriptional bursting, the underlying mechanisms remain unclear (Zhang et al., 2024; Fukaya et al., 2016). Second, Pol II molecules transiently form clusters at promoters to initiate transcription, but these clusters dissipate rapidly, even as transcription continues (Ryu et al., 2024; Pancholi et al., 2021; Cho et al., 2016; Cisse et al., 2013). The origin of additional Pol II molecules required for ongoing transcription is not yet understood. Third, transcription is not restricted to promoters; it also occurs at enhancer regions and intervening sequences, producing short, unstable, non-coding transcripts known as enhancer RNAs (eRNAs) (Qiu et al., 2026; Carullo et al., 2020; De Santa et al., 2010; Kim et al., 2010; Ashe et al., 1997; Tuan et al., 1992). Several studies have shown that eRNA transcription can influence gene activity (Racko et al., 2018; Kaikkonen et al., 2013; Li et al., 2013), yet the precise mechanisms remain unresolved (Wang et al., 2026; Qiu et al., 2026). It has been proposed that either the process of eRNA transcription or the eRNA molecules themselves — or both — may mediate transcriptional regulation (Qiu et al., 2026; Natoli & Andrau, 2012). Fourth, gene promoters not only loop to interact with enhancers, as described in the classical looping model (Wendt et al., 2008; Splinter et al., 2006), but also scan and bind to other DNA sequences in the vicinity and beyond, potentially within dynamic transcriptional condensates (Ryu et al., 2024; Pancholi et al., 2021). How these observations integrate into a unified model of transcriptional regulation is still unclear. Transcriptional regulation is highly dynamic, especially during development (Levine & Davidson, 2005), when gene expression patterns change rapidly to guide cell fate decisions. This dynamic nature makes developmental systems an excellent model for studying the principles of transcriptional control.

Several strategies have been employed to investigate the role of eRNA transcription in gene regulation. Genomics-based methods such as global run-on sequencing (GRO-seq), precision nuclear run-on sequencing (PRO-seq), and cap analysis gene expression (CAGE) have enabled genome-wide analysis of nascent transcripts (Carullo et al., 2020; Lewis et al., 2019). However, these bulk assays average the behavior of millions of cells, limiting their ability to resolve the spatiotemporal dynamics of eRNA transcription. RNA interference techniques—including short hairpin RNAs (shRNAs), small interfering RNAs (siRNAs), and locked nucleic acids (LNAs)—have been used to knock down eRNA transcripts in cell culture, providing insight into their cellular roles (Pnueli et al., 2015; Hsieh et al., 2014; Melo et al., 2013). Nonetheless, cell culture systems may not fully capture the complexity of eRNA function during embryonic development. In contrast, fluorescent RNA *in situ* hybridization chain reaction (FISH HCR) enables the visualization of eRNA transcriptional dynamics and their relationship to gene transcripts at the single-cell level and was shown in cell culture (Shibayama et al., 2017).

In this study, we implemented an imaging-based approach using FISH to investigate the characteristics of eRNA transcription during development. We selected the early patterning of the anterior-posterior (AP) axis as a model system, focusing on the well-characterized AP patterning genes of the fruit fly *Drosophila melanogaster* (*Drosophila*) (Schroeder et al., 2011; Schroeder et al., 2004). This system offers several advantages: (1) a wealth of characterized patterning enhancers, and (2) the blastoderm embryo’s single layer of nuclei, which facilitates imaging at single-cell resolution.

We concentrated our analysis on enhancers of gap and pair-rule genes, which respectively define the early anterior–posterior axis and the periodic segmentation pattern in *Drosophila* embryos. Using precisely staged embryos, we investigated the endogenous dynamics of eRNA expression at native genomic loci. To further dissect eRNA activity, we developed an enhancer reporter system that allowed controlled assessment of eRNA transcription—both in fixed embryos via in situ HCR and in live embryos using an MS2 reporter assay. Our results show that eRNAs are transcribed not only from active enhancer regions but also from adjacent sequences, even when these are replaced by foreign bacterial sequences. Notably, eRNA transcription displays spatial and temporal patterns that closely parallel those of their associated genes.

## Results

### Characterization of eRNA Transcriptional Properties in the Endogenous Gene Locus

To investigate the properties of eRNA transcription, we focused on the early anterior-posterior (AP) patterning genes *Krüppel* (*Kr*) and *giant* (*gt*). For *Kr*, we analyzed the enhancers KrCD1 (−3,948 bp to -2,435 bp upstream of the *Kr* transcription start) and KrCD2 (−2,605 bp to -121), both of which are active in a central domain along the embryo’s AP axis (El-Sherif & Levine, 2016; Hoch et al., 1990) (**Figure 1A’**). For *gt*, we examined the enhancers gt(−1) [-1,284 bp to -45 bp upstream of the *gt* transcription start] and gt(−3) [-2,500 bp to -1,291 bp]: gt(−1) is active in both, anterior and posterior domains, whereas gt(−3) activity is restricted to the posterior domain (Schroeder et al., 2004) (**Figure 1B’**). All analyses were performed in *Drosophila* embryos at nuclear cycle 14 (NC14), corresponding to blastoderm cellularization approximately 2 hours 10 minutes to 3 hours after egg laying (Foe et al., 1993).

**Figure 1.**
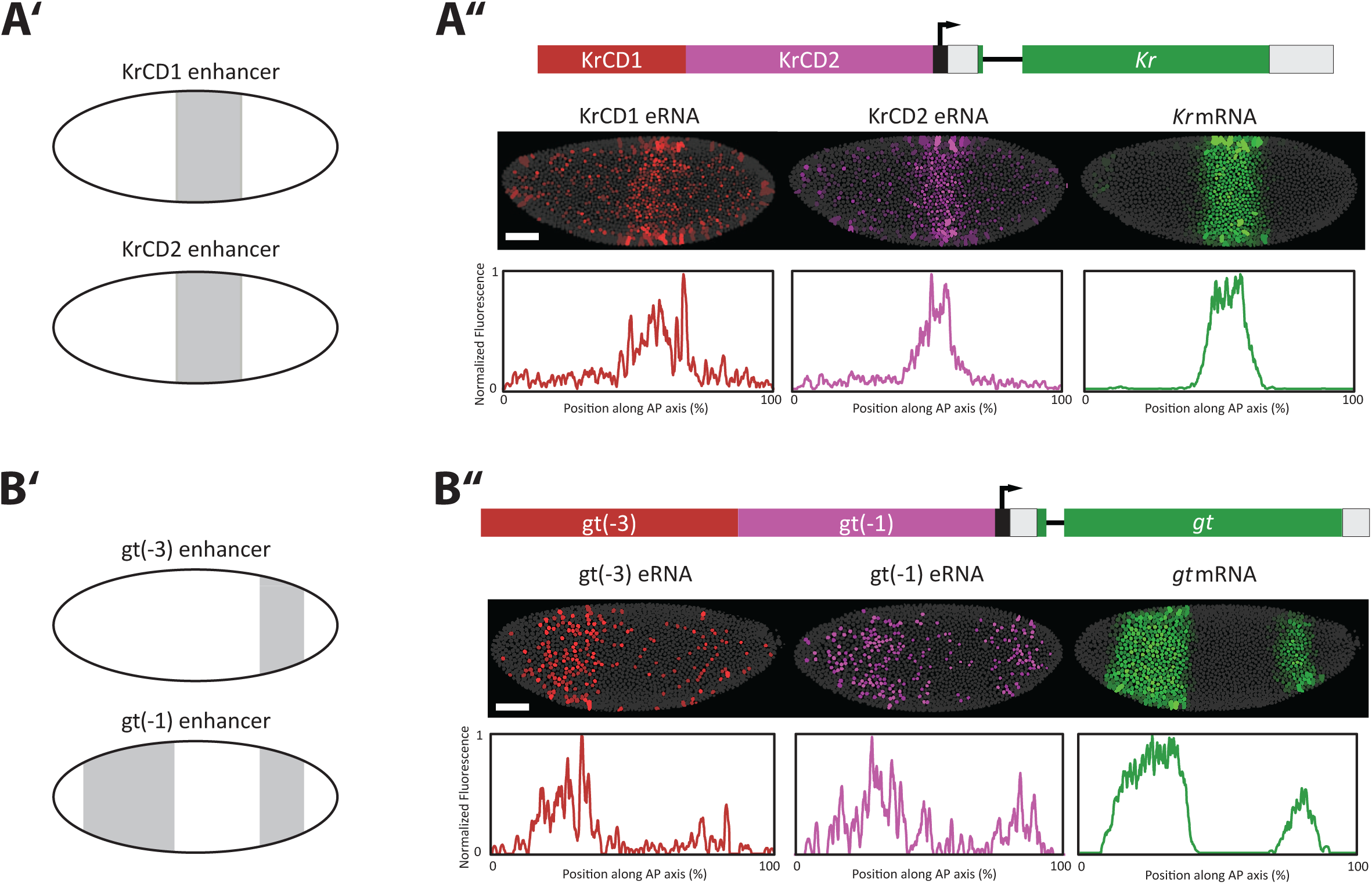
Spatial expression of eRNAs from gap gene enhancers. *Kr* locus and *gt* locus are shown. (**A′, B′**) Schematics illustrate the activity patterns of selected enhancer regions in the blastoderm embryo. (**A′**) The KrCD1 and KrCD2 enhancers drive expression in the central domain. (**B′**) The gt(−3) enhancer is active in the posterior domain, whereas the gt(−1) enhancer drives expression in both anterior and posterior domains. (**A″, B″**) eRNA (red and purple) and mRNA (green) transcripts were visualized together in the endogenous gap gene locus in whole *Drosophila* embryos at nuclear cycle 14 (NC14) using *in situ* HCR staining. Nuclei are shown in grey. Fluorescence signals along the dorsal–ventral axis were summed and normalized to the maximum intensity to generate eRNA distribution profiles along the anterior–posterior (AP) axis (position in %). (**A″**) eRNA transcripts from the KrCD1 and KrCD2 enhancers are detected in the central AP domain, overlapping with the *Kr* mRNA expression region. (*B″*) *gt* mRNA forms anterior and posterior domains, with eRNA from the gt(−3) and gt(−1) enhancers detected at similar AP positions. Anterior is left, dorsal is up. Scale bars: 50 µm.

To visualize the low-abundance eRNA transcripts, we employed high-sensitivity fluorescence *in situ* hybridization chain reaction (HCR) staining (Choi et al., 2018). This method enables multiplexed detection (up to five RNA targets simultaneously), quantitative analysis at single-molecule resolution, and sensitive identification of low-abundance RNAs through automatic background suppression. Here, we stained one mRNA and two eRNAs in the same embryo, allowing us to precisely study their relationship. HCR staining was combined with high-resolution confocal microscopy. For HCR staining, probes were uniformly distributed over the assessed region, covering about 50 bp per probe (for probe LOT numbers see **Supplementary Table 2**).

Consistent with previous reports, eRNA expression was predominantly nuclear. For the KrCD1 and KrCD2 enhancers, eRNA transcripts were detected in a central domain of the embryo, mirroring the expression pattern of *Kr* mRNA (**Figure 1A’’**). Similarly, eRNA transcripts from gt(−1) and gt(−3) enhancers were enriched in anterior and posterior domains, closely resembling the *gt* mRNA expression pattern (**Figure 1B’’**). Notably, gt(−3) eRNA transcripts were strongly expressed in the anterior domain, despite enhancer activity being confined to the posterior (Schroeder et al., 2004).

These observations demonstrate that eRNA transcription is spatially correlated with mRNA expression of the corresponding gene. The activity of gt(−3) enhancer eRNA transcription at the anterior was surprising in contrast to the reported gt(−3) activity at the posterior, raising questions about the broader landscape of eRNA transcription within gene loci.

### Profiling the Landscape of eRNA Transcription within the Gene Locus

Given the observed spatial correlation between eRNA and mRNA transcription, we next sought to systematically profile the abundance of eRNA transcription across a gene locus. Two possible configurations might be expected: (i) eRNA transcripts may be uniformly expressed throughout the gene locus, or (ii) eRNA transcripts may arise from discrete, defined regions within the locus.

To this end, we analyzed the 6 kb upstream region of the *Kr* locus, which encompasses the KrCD1 and KrCD2 enhancers. For *in situ* HCR staining, we designed detection windows of 500 bp to map eRNA transcriptional activity at high resolution. As previous studies have demonstrated that eRNAs are frequently bidirectionally transcribed (Hah et al., 2013; Melgar et al., 2011), we probed for both, sense and antisense eRNA transcripts. Sense and antisense probes were alternating for detection of sense and antisense eRNA transcripts in the same embryo (4-5 probes per detection window for sense and antisense, respectively).

eRNA distribution was analyzed at nuclear cycle 13 (NC13; syncytial blastoderm, approximately 1 hour 50 minutes to 2 hours 10 minutes after egg laying [Blythe & Wieschaus, 2015; Foe et al., 1993]), a developmental stage at which both, mRNA and eRNA transcriptional foci, are readily detectable.

High-resolution confocal imaging was used to detect *Kr* mRNA and eRNA transcription foci. The intensity of eRNA foci overlapping with *Kr* mRNA foci was analyzed along the 6kb region. Our high-resolution mapping revealed hotspots of eRNA transcription for both, sense and antisense probes (**Figure 2**, arrow heads). We identified several prominent peaks of eRNA transcription: one upstream of the KrCD1 enhancer (sense and antisense eRNA signal), one in the KrCD1 enhancer (sense eRNA signal), and a third one in the middle of the KrCD2 enhancer (sense and antisense eRNA signal) suggesting that these regions serve as sources of eRNA production within the *Kr* locus. In addition, distinct regions within the KrCD1 and KrCD2 enhancers exhibited local minima in eRNA signal.

**Figure 2.**
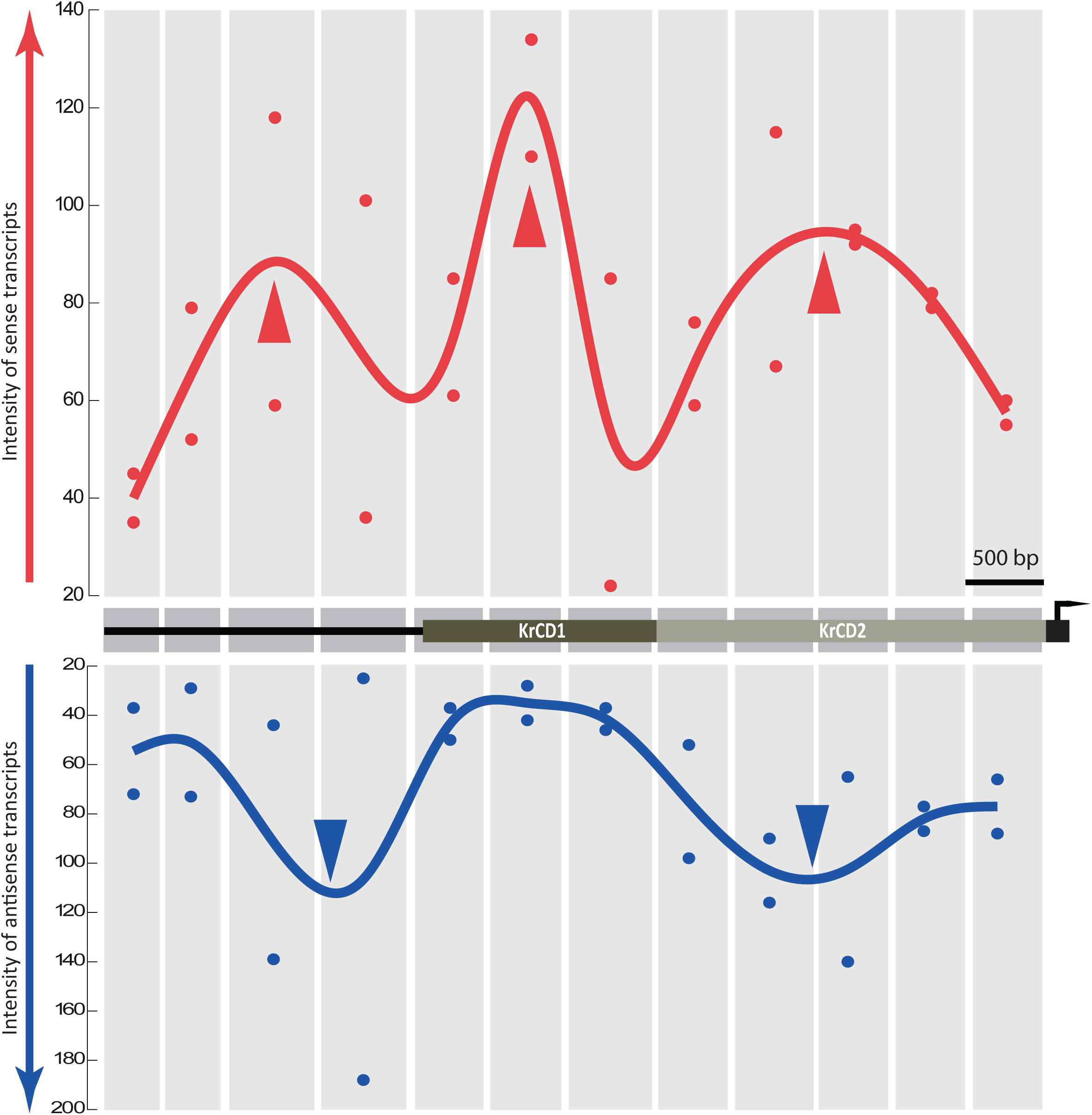
High resolution mapping of eRNA transcription along the upstream sequence of the endogenous *Kr* locus. A 6-kb upstream region of the endogenous *Kr* locus containing KrCD1 and KrCD2 enhancers was analyzed. eRNA transcripts were visualized by *in situ* HCR staining at NC13. Sense (red) and anti-sense (blue) eRNA transcripts were detected using 4-5 probes spanning ∼500 bp windows. Two embryos were imaged per window. High-resolution confocal imaging was used to detect *Kr* mRNA and eRNA transcription foci. The y-axis shows the intensity of eRNA foci overlapping with *Kr* mRNA foci. Three maxima of sense eRNA - one upstream of KrCD1, one in KrCD1, and one in KrCD2 - and two maxima of anti-sense eRNA – one upstream of KrCD1 and one in KrCD2 – were observed (arrow heads).

Collectively, these results indicate that eRNA transcription is not uniformly distributed but instead arises from discrete hotspots within the gene locus. The spatial organization of these hotspots may be shaped by the occupancy of transcription factors and the transcriptional machinery at enhancers and promoters.

### Establishing an Enhancer Reporter System to Study eRNA Transcriptional Dynamics

Our analysis of eRNA transcription at endogenous gene loci revealed that eRNA synthesis in the embryo occurs in spatial patterns closely mirroring those of mRNA expression. However, in the endogenous context, eRNA transcription may be influenced by the entire regulatory landscape of the genomic locus, which complicates the interpretation of individual enhancer activities. To overcome this limitation, we developed a *Drosophila* enhancer reporter system that enables visualization of eRNA transcription in a controlled and reproducible setting outside of the endogenous locus.

This reporter system is based on the pbPHi vector, which contains the *Drosophila yellow* reporter gene with MS2 stem loops inserted into its 5′ untranslated region (UTR) and an attB site for site-specific genomic integration (Bothma et al., 2014; Perry et al., 2010). We cloned gap gene or pair-rule gene enhancers, along with their corresponding endogenous promoter sequences, into this construct to assay eRNA transcription (**Figure 3A**). This system allows for the analysis of eRNA and *yellow* mRNA expression dynamics in fixed embryos using FISH, as well as in live embryos via the MS2-MCP system (Garcia et al., 2013). The MS2-MCP system enables real-time visualization of nascent transcription: male flies carrying the enhancer>MS2-yellow construct are crossed with females expressing a ubiquitously driven MS2 coat protein (MCP)-GFP fusion. In cells where the enhancer is active, the MS2-yellow transcript is produced and bound by MCP-GFP, resulting in localized GFP enrichment at sites of active transcription.

**Figure 3.**
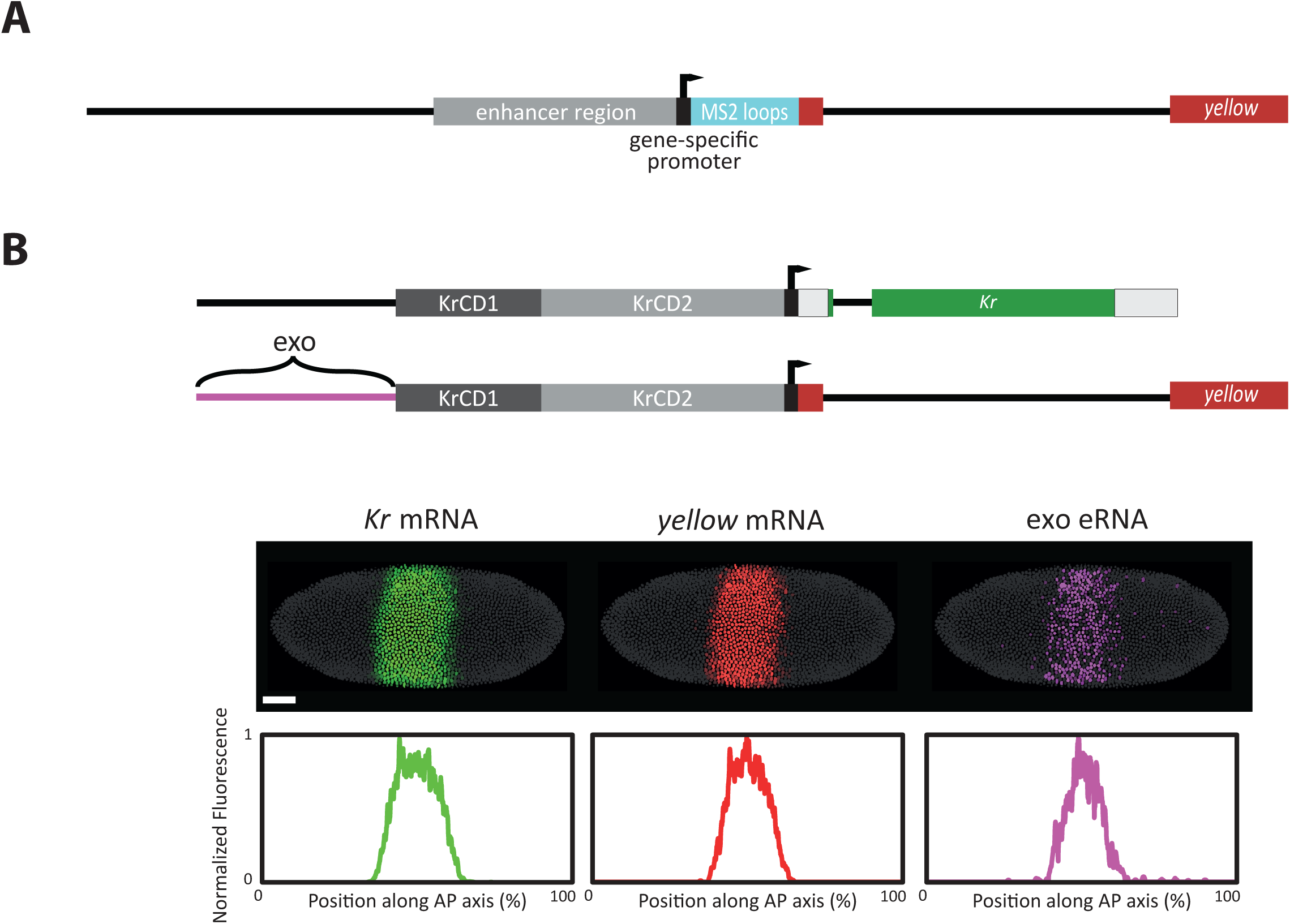
eRNA activity extends to reporter gene vector sequences. (**A**) Schematic of the enhancer reporter construct: enhancer regions were cloned into the pbPHi plasmid upstream of gene-specific promoters. All constructs contain the *yellow* gene with 24xMS2 stem loops for live imaging, and the mini-white marker for selection. Site-specific integration was achieved via PhiC31 integrase. (**B**) Transcriptional profile of the *Kr* enhancer reporter containing KrCD1 and KrCD2 enhancers: *In situ* HCR staining of whole embryos (NC14) visualized *Kr* mRNA, *yellow* mRNA, and a bacterial exogenous (exo) sense eRNA. Anterior is left, dorsal is up. Scale bars: 50 µm.

Using PhiC31 integrase-mediated site-specific transgenesis, we generated several transgenic *Drosophila* lines carrying enhancer region>MS2-yellow constructs. To validate this system, we utilized the previously characterized 6 kb upstream sequence of the *Kr* locus containing the KrCD1 and KrCD2 enhancers (KrCD1+2 reporter line). To distinguish eRNA transcripts generated by the transgenic construct from those produced at the endogenous *Kr* locus, we designed *in situ* HCR probes targeting an *Escherichia coli* (*E. coli*) derived exogenous (exo) sequence integrated with the reporter construct. Notably, also in this situation, eRNA’s were found upstream of the KrCD1 enhancer, in the *E. coli* derived exo vector sequences, and they spatially correlated with mRNA transcripts from the KrCD1+2 reporter (**Figure 3B**) as well. This shows that eRNAs can be generated in foreign bacterial sequences when these are brought close to an active enhancer. Thus, the eRNAs upstream of the KrCD1 enhancer (**Figure 2**) likely were induced by these enhancers, and were not due to previously overlooked “weak” enhancers in the endogenous gene.

### Analysis of eRNA Transcription in the Absence of a Gene Promoter

To determine whether eRNA transcription merely reflects background transcriptional noise at the gene promoter, we engineered an enhancer reporter system utilizing the stripe-specific *even-skipped* (*eve*) enhancer 1+5 (+6,616 bp to +8,210 bp of the *eve* transcription start), which drives expression in stripes 1 and 5 of the characteristic seven-striped *eve* gene pattern (Fujioka et al., 1999). Notably, this enhancer does not produce detectable eRNA at the endogenous *eve* locus (**Supplementary Figure 1A**), enabling us to analyze eRNA transcripts originating exclusively from the reporter construct without interference from endogenous eRNA.

Consistent with findings from the KrCD1+2 enhancer reporter, the eve1+5 reporter also exhibited eRNA expression from vector sequences adjacent to the stripe enhancers, and also these exo eRNAs coincided spatially with mRNA expression of the *yellow* reporter gene (**Figure 4A**). While the *eve* enhancer 1+5 shows no detectable eRNA expression at the endogenous locus, eRNA transcription is observed in the reporter construct, which may be enabled or enhanced by as yet unknown features of the reporter construct.

**Figure 4.**
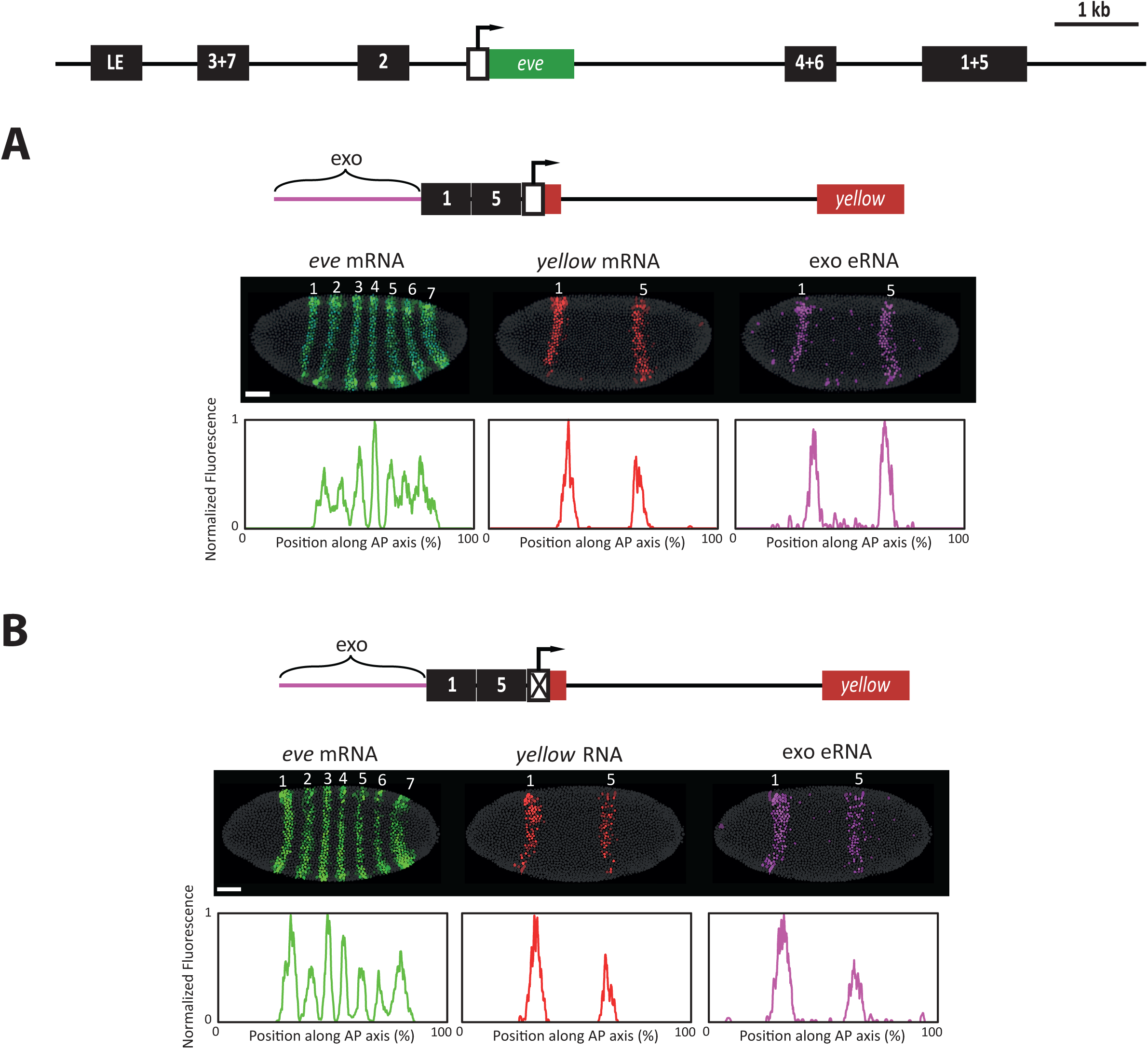
eRNA expression in the presence and absence of a promoter. *In situ* HCR staining (NC14) visualized mRNA (green/red) and sense eRNA (purple) together in whole embryos. Nuclei are gray. Fluorescence along the dorsal–ventral axis was integrated and normalized to generate AP distribution plots. (**A**) eve1&5 enhancer reporter: the stripe-specific *eve* enhancer 1+5 drives *yellow* mRNA and eRNA at stripes 1 and 5. Anterior is left, dorsal is up. (**B**) eve1&5wop enhancer reporter (wop: without *eve* promoter [-50 bp to +50 bp of the *eve* transcription start])): Also when the promoter is deleted, the same enhancer drives weaker *yellow* RNA and eRNA at stripes 1 and 5 independent from a promoter. *yellow* RNA transcripts are nuclear localized resembling eRNA transcripts. Anterior is left, dorsal is up. Scale bar: 50 µm.

In addition, we generated a modified enhancer reporter lacking the gene promoter (eve1+5wop; wop: without *eve* promoter, i.e. lacking the interval from -50 bp to +50 bp relative to the *eve* transcription start site). In this line, RNA transcription was still detected, indicating that eRNA transcription can occur independently of the gene promoter. Both, exo and *yellow* region transcripts, were observed (**Figure 4B**). In the absence of a gene promoter, *yellow* transcripts lost their typical mRNA features and instead exhibited nuclear localization resembling eRNA features. Cytoplasmic expression was minimal, as is typical for other eRNA transcripts.

In summary, these findings demonstrate that eRNA transcription can occur independently of promoter-driven transcription. As next step, we tested if eRNA activation by enhancers can be blocked by insulators.

### Analyzing the Influence of Insulators on eRNA Transcription

To test the influence of insulators on eRNA transcription, we established a 3&7&2 enhancer reporter line. This construct contains a 3.6 kb region upstream of the *eve* transcription start site, which includes the stripe-specific *eve* enhancers 3+7 (−3,817 bp to -3,307 bp relative to the *eve* transcription start site) and 2 (−1,557 bp to -1,073 bp). The 3+7 enhancer drives expression at stripes 3 and 7 in the characteristic *eve* pattern (Small et al., 1996), while the enhancer 2 drives expression at stripe 2 (Small & Levine, 1991). As these enhancers do not produce detectable eRNA at the endogenous *eve* locus (**Supplementary Figure 1B+C**), our system enables the analysis of eRNA transcription in a controlled context, free from endogenous eRNA. We visualized both, mRNA and sense eRNA expression, using *in situ* HCR staining.

The 3&7&2 enhancer reporter line exhibited eRNA expression at stripe positions 2, 3, and 7, coinciding with the *yellow* mRNA expression pattern (**Figure 5A**). In line with the observation for eRNAs derived from the gt(−3) enhancer (**Figure 1B’’**, gt(−3) eRNA), eve enhancer 2 showed eRNA expression at stripe positions 2,3, and 7, not only at stripe position 2 (**Figure 5A**, Str2 eRNA).

**Figure 5.**
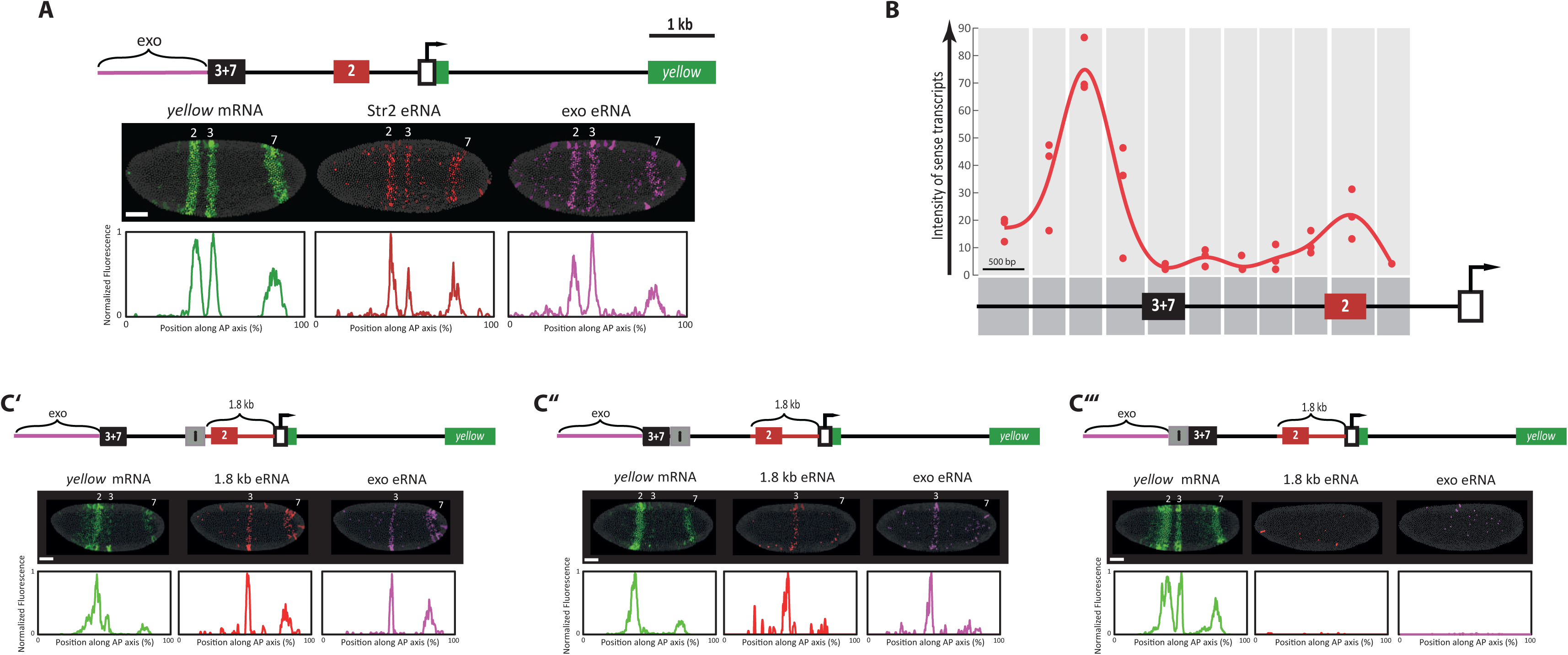
Influence of insulators on eRNA transcription. *In situ* HCR staining (NC14) visualized together mRNA (green) and sense eRNA (red/purple) in whole embryos. (**A**) eve3&7&2 enhancer reporter: *yellow* mRNA, Str2 eRNA, and exo eRNA are detected at stripes 2, 3, and 7. (**B**) Transcriptional profiling: A 5.5-kb upstream region (including endogenous enhancers and exogenous pbPHi sequence) was analyzed in 500-bp windows. The number of eRNA foci overlapping with *yellow* mRNA foci is plotted per embryo (n=3). A maximum of eRNA is detected in the exogenous region, a small maximum just downstream of *eve* enhancer 3+7, and another one at *eve* enhancer 2. (**C’-C’’’**) eRNA transcription blocked by su(Hw) insulator configurations: (**C’**) 3&7-su(Hw) upstream-2 transgene: Insulator is located 1.8 kb upstream of the *eve* transcription start site. *Yellow* mRNA is detected at stripe 2 and weakly at 3/7; eRNAs at stripes 3 and 7 only. (**C’’**) 3&7-downstream su(Hw)-2 transgene: Insulator is positioned just downstream of enhancer 3+7. *Yellow* mRNA detected at stripes 2 and 7; eRNA at stripe 3 and weakly at 7. (**C’’’**) su(Hw)-3&7&2 transgene: Insulator was inserted upstream of enhancer 3+7. *Yellow* mRNA is expressed at stripes 2, 3, and 7; no eRNAs are detected. Anterior is left, dorsal is up. Scale bars: 50 µm.

Profiling the transcriptional landscape revealed maxima of eRNA expression at the exo region, a small maximum just downstream of *eve* enhancer 3+7, and another one at *eve* enhancer 2 (**Figure 5B**). Reminiscent of our observations at the endogenous *Kr* locus, local minima of eRNA transcription were detected at enhancer region 3+7 and near the promoter. Since the profile lacks a sharp rise followed by a gradual decline and instead displays multiple maxima and minima, the existence of a cryptic promoter can be ruled out.

The global maximum at the exo region appears to represent a region of strong transcriptional activity responsible for eRNA synthesis. To test this, we inserted an insulator sequence at various positions within the 3&7&2 enhancer reporter construct to investigate the effect of insulators on eRNA transcription, while the unaffected enhancer region 2 served as internal control. We utilized a 347 bp fragment containing su(Hw) binding sites (hereafter “su(Hw) insulator”), derived from the gypsy retrotransposon (Geyer & Corces, 1992), which is well characterized for its ability to block enhancer–promoter communication in *Drosophila* (Cai & Levine, 1995). Three insulator-containing reporter lines were generated: (i) su(Hw) insulator upstream of enhancer 2 (3&7-su(Hw) upstream-2), (ii) su(Hw) insulator downstream of enhancer 3+7 (3&7-downstream su(Hw)-2), and (iii) su(Hw) insulator upstream of enhancer 3+7 (su(Hw)-3&7&2). We then quantified mRNA transcripts from the exonic *yellow* region and eRNA transcripts from either the exo region or the 1.8 kb region upstream of the *eve* transcription start site.

In the 3&7-su(Hw) upstream-2 line, *yellow* mRNA was present at stripe 2 but only weakly at stripes 3 and 7. Residual mRNA at stripes 3 and 7 may result from incomplete insulation or from Pol II run-through, as these transcripts were predominantly nuclear and eRNA-like. Notably, *yellow* mRNA pattern at stripe 2 position was slightly weaker and not as concise as without the insulator (compare **Figure 5C’** to **Figure 5A**, *yellow* mRNA) eRNA transcription was detected only at stripes 3 and 7, revealing the insulator’s ability to block eRNA transcription (**Figure 5C’**), but seems to work different than with mRNA, since stripe 2 effect is blocked not only for exo eRNA but also for 1.8 kb eRNA.

Similarly, in the 3&7-downstream su(Hw)-2 line, eRNA transcription was blocked at stripe 2 (**Figure 5C’’**). At stripe 7, eRNA expression was reduced while *yellow* mRNA expression increased, consistent with previous findings that *eve* expression at stripe 7 is driven not only by enhancer 3+7 but also by the intervening sequence between enhancers 3+7 and 2 (Small et al., 1996).

In the su(Hw)-3&7&2 line, eRNA transcription was completely abolished, while *yellow* mRNA was unaffected and expressed at stripes 2, 3, and 7 (**Figure 5C’’’**).

This shows that the spatial pattern of exo eRNAs is regulated by enhancers and can be blocked by insulators in a manner similar to enhancer–promoter interactions. However, the absence of eRNA transcripts at stripe position 2 for the 1.8 kb eRNA stainings remains unexplained, as this region should not be affected by the insulator—given that the *eve* enhancer 2 is not separated from the 1.8 kb region by it. One possible explanation is that the su(Hw) insulator exerts two distinct effects: (1) enhancer insulation, analogous to its function in blocking promoter–enhancer communication, and (2) a stripe 2–specific silencing effect that operates in a position-independent manner.

Collectively, these insulator studies provide strong evidence that eRNA transcription is independently driven of gene promoters. This raises the question of whether, and if so, how eRNA transcription influences gene transcription itself. Our novel experimental approach could be used to address this question by leveraging the MS2-MCP live imaging capabilities of our enhancer reporter system.

### Promoter Burst Characteristics is Influenced by the Interaction of Insulator and Stripe 2

To investigate the influence of the insulator on stripe 2 activity, comparative live imaging was performed on the 3&7&2 and the 3&7-su(Hw) upstream-2 enhancer reporter lines from early to late NC 14, using two-minute time intervals. The analysis focused specifically on stripe position 2, where mRNA expression is affected by a nearby insulator in an unexpected manner (**Figure 5C’**, *yellow* mRNA slightly weaker and less concise).

In the 3&7&2 reporter, prominent GFP puncta were evident at stripe position 2, signifying active transcription sites that mirrored the *yellow* mRNA expression pattern seen in HCR stainings (**Movie S1**, **Figure 6A**). In the 3&7-su(Hw) upstream-2 line, where the insulator blocks eRNA expression at stripe position 2 and partially inhibits mRNA expression (**Figure 5C’**), GFP puncta were also visible, but slightly weaker (**Movie S2**, **Figure 6B**).

**Figure 6.**
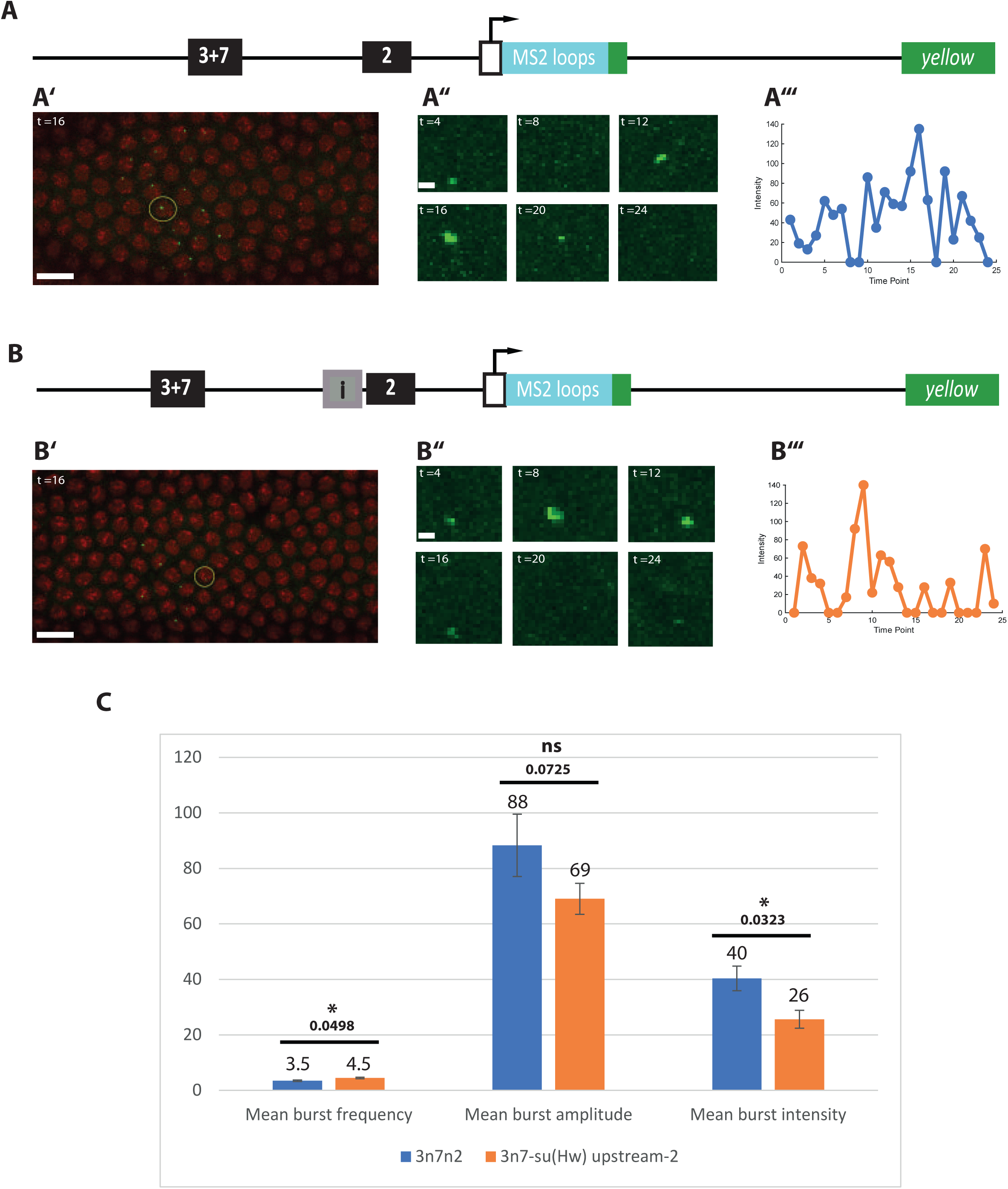
Analysis of the influence of an insulator on transcriptional burst activity at *eve* stripe 2. (**A**, **B**) Live imaging of embryos expressing “3&7&2>MS2-*yellow*; MCP-GFP; Histone-RFP” (A) and “3&7-su(Hw) upstream-2>MS2-*yellow*; MCP-GFP; Histone-RFP” (B) from early to late NC 14 provides time-lapse visualization (2-minute intervals) of active transcriptional sites, with nuclei marked in red (Histone-RFP) and nascent transcripts visualized as green GFP puncta. Yellow circles indicate analyzed nuclei, and anterior is to the left (scale bars: 10 µm). For each condition, representative nuclei traces show transcriptional bursting over time (A’’, B’’: puncta snapshots, scale bars: 1 µm; A’’’, B’’’: intensity time course). (**C**) Quantitative comparison reveals a significant increase in mean burst frequency (bursts were identified as periods when the signal exceeds 5% of the maximum trace intensity, 4.5 ± 0.25 vs. 3.5 ± 0.25, p = 0.0498) and a significant decrease in mean burst intensity (average signal across the entire trace, 26 ± 3.24 vs. 40 ± 4.45, p = 0.0323), but no significant difference in mean burst amplitude (average of the maximum signal intensities within each burst, 69 ± 5.58 vs. 88 ± 11.20, p = 0.0725) in the 3&7-su(Hw) upstream-2 reporter relative to 3&7&2 (n=4 nuclei in one embryo per reporter line).

Live imaging analysis of 4 nuclei in one embryo per reporter line revealed a statistically significant increase in mean transcriptional burst frequency, where bursts were identified as periods when the signal exceeds 5% of the maximum trace intensity, in the 3&7-su(Hw) upstream-2 line (4.5 ± 0.2450) compared to the 3&7&2 line (3.5 ± 0.2500; p = 0.0498). The mean burst amplitude (average of the maximum signal intensities within each burst), while slightly reduced in the 3&7-su(Hw) upstream-2 line (69 ± 5.58) versus the 3&7&2 line (88 ± 11.20), did not reach statistical significance (p = 0.0725). Importantly, mean burst intensity (average signal across the entire trace) was significantly lower in the 3&7-su(Hw) upstream-2 line (26 ± 3.24) relative to the 3&7&2 reporter (40 ± 4.45; p = 0.0323) (**Figure 6C**).

These subtle shifts could reflect direct insulator effects on chromatin architecture or local Pol II availability, indirect consequences of blocked eRNA transcription, or a combination of both. While eRNA suppression might deplete a local Pol II pool and prompt compensatory increases in burst frequency, the modest magnitude of changes precludes definitive attribution to any single mechanism.

## Discussion

Enhancer RNAs (eRNAs) have been identified across a broad range of organisms — including *Drosophila*, *Caenorhabditis elegans*, mice, and humans (Meers et al., 2018; Chen et al., 2013; Kim et al., 2010; Ashe et al., 1997) — highlighting their potential as important regulators of transcription. In this study, we established an imaging-based framework to investigate eRNA transcriptional dynamics at the single-cell level in a controlled system. By integrating high-sensitivity HCR staining, high-resolution confocal microscopy, and enhancer reporter assays, we elucidated the characteristics of eRNA transcription in whole fixed or live *Drosophila* embryos.

Our findings provide new insights into the spatial and temporal dynamics of eRNAs and their functional relation to enhancers. We observed that eRNAs are expressed in similar temporal and spatial patterns as their associated enhancers (**Figure 1**). While temporal co-activity between eRNAs and enhancers had already been reported (Gao et al., 2023; Mikhaylichenko et al., 2018), the overlap in spatial expression patterns has so far been largely assumed. Our data therefore strengthen this link and support the idea that eRNAs can serve as reliable reporters of enhancer activity in both time and space.

Interestingly, we detected eRNA transcription not only within canonical enhancer regions but also in nearby sequences (**Figure 2**), challenging the prevailing view that eRNAs originate strictly from enhancer regions (Franco et al., 2018; Cheng et al., 2015). Instead, our results indicate that eRNAs can be produced in the vicinity of active enhancers. Critically, this eRNA production occurs independently of any promoter presence (**Figure 4**), underscoring that enhancers alone are sufficient to drive non-coding transcription in their vicinity.

eRNA transcriptional activity can be effectively blocked by the introduction of an insulator element (**Figure 5C**), suggesting that local chromatin architecture plays a critical role in defining the boundaries of eRNA activity. Moreover, eRNA transcription that can be blocked by insulators suggests that the sources of eRNA transcription may possess promoter-like characteristics. This idea aligns with previous studies showing that many active enhancers initiate local transcription through factors and mechanisms similar to those used by canonical promoters (Tippens et al., 2018). Genome-wide studies using techniques such as CAGE and GRO-seq have revealed that enhancers and promoters share similar chromatin marks (e.g., H3K27ac, H3K4me1/3), transcription factor occupancy, and RNA Pol II recruitment, and that both can serve as sites of divergent transcription initiation (Kristjánsdóttir et al., 2020, Tippens et al., 2018).

Even more strikingly, we observed eRNA production in unrelated bacterial sequences when these were positioned near active enhancers (**Figure 3B**). This finding indicates that eRNA transcription is not limited to endogenous enhancer-associated sequences and can be induced simply by proximity to an active enhancer. Therefore, the presence of eRNAs in such regions cannot be explained by overlooked weak enhancers in the endogenous genome but rather reflects a broader property of enhancer-driven transcription.

We also identified a potential insulator effect on a downstream enhancer that could arise through blocking eRNAs transcribed from upstream regions (**Figure 6**). This observation raises the intriguing possibility that eRNAs themselves may contribute to enhancer interference or insulation effects, suggesting a more complex and dynamic regulatory interplay between neighbouring enhancers.

We hypothesize that RNA Polymerase II (Pol II) clusters arrive at the gene locus in an episodic manner. Upon arrival, Pol II molecules are distributed in the vicinity of active enhancers. Regions of eRNA transcription might act as reservoirs, storing Pol II around the gene locus and facilitating its recycling to the gene promoter when needed, thereby supporting sustained mRNA transcription. This model suggests that transcriptional bursting is modulated by the frequency of Pol II cluster recruitment. Nevertheless, more in detail analysis of live imaging constructs with other enhancers and insulators has to be performed to develop a universally valid model of the role of eRNA transcription in gene regulation.

Despite these observations, several questions remain. The precise molecular mechanisms by which eRNA transcription modulates Pol II dynamics and/or chromatin architecture are still unresolved. Additionally, while our findings hint at the significance of eRNA transcription during early embryonic development, it is important to determine whether similar mechanisms operate in other tissues and developmental stages.

Finally, our reporter vector system proved highly useful for dissecting enhancer–eRNA interactions and could be adapted for wider applications. In particular, it provides a versatile tool to experimentally test enhancer activity, insulator mechanisms, and noncoding transcription in diverse genomic contexts.

## Availability of Data and Materials

Data and materials are available upon request from the authors.

Code is available in **Supplementary File 1**. Program 1 was used to detect transcription sites in **Figure 1A’’+1B’’**, **Figure 2**, **Figure 3B**, **Figure 4**, and **Figure 5**. Program 2 was used to create AP and DV plots in **Figure 1A’’+1B’’**, **Figure 3B**, **Figure 4**, and **Figure 5A+5C’-C’’’**. Program 3 was used to generate eRNA transcription profiles in **Figure 2** and **Figure 5B**. Program 4 was used for burst analysis in **Figure 6**.

## Acknowledgement

This work is supported by a DFG grant (EL 870/2-1) to EE, and a doctoral fellowship from the German Academic Scholarship Foundation to CM.

## Material and Methods

### Fly Culture

*Drosophila melanogaster* were maintained in standard vials containing *Drosophila* food prepared as follows (for 10 L): 12.5 L water, 53.2 g agar, 900 g cornmeal, 600 g sugar beet syrup, 150 g dry yeast, 30 g Nipagin, 120 mL 70% ethanol, and 50 mL propionic acid. Flies were kept at 18 °C for stock maintenance. For mating and egg collection, flies were transferred to 25 °C.

### Enhancer Reporter Constructs

All enhancer reporter constructs were generated using the pbPHi plasmid, which contains a mini-white marker, a minimal *eve* promoter (−50 bp to +50 bp relative to the *eve* transcription start site), a 24xMS2-*yellow* cassette, and an attB site (Bothma et al., 2014). Enhancer regions (Fujioka et al., 1999; Small et al., 1996; Fujioka et al., 1995; Small & Levine, 1991), the su(Hw) insulator (Cai & Levine, 1995; Geyer & Corces, 1992), intergenic regions, and pbPHi backbone regions were PCR-amplified and inserted into the linearized pbPHi plasmid using In-Fusion cloning (Takara Bio Europe AB). Plasmid linearization was performed with NotI or NotI+BamHI to excise the minimal *eve* promoter as required. Primer sequences are listed in **Supplementary Table 1**.

### *Drosophila* Transgenesis

Enhancer reporter constructs were integrated into chromosome III of the Vk33 strain (RRID:BDSC_9750) via PhiC31 integrase-mediated site-specific transgenesis (BestGene Inc.). Transgenic flies were identified by the mini-white marker and crossed to establish homozygous lines for each enhancer reporter construct.

### Egg Collections

For embryo collections, adult flies were placed in cages with agar apple juice plates (2.375% w/v agar, 2.5% w/v sucrose, 25% v/v apple juice) and kept at 25 °C for the desired collection period.

### *In Situ* Hybridization and Imaging of Fixed Embryos

*In situ* hybridization was performed using the third-generation hybridization chain reaction (HCR v3.0; Choi et al., 2018). Probe sets and hairpins were obtained from Molecular Instruments (see **Supplementary Table 2** for lot numbers). Imaging was conducted using a Leica TCS SP5 II confocal laser scanning microscope with 20x or 63x objectives at 2048 × 1024 resolution.

### Live Imaging

MCP-GFP and Histone-RFP expressing females (Garcia et al., 2013) were crossed with males carrying either the 3&7&2 enhancer>MS2-*yellow* or 3&7-su(Hw) upstream-2 enhancer>MS2-*yellow* constructs. Eggs were collected over 1 hour and incubated for an additional 30 minutes at 25 °C. Embryos were dechorionated in bleach for 1 min 40 sec, mounted using the hanging drop method, and covered with halocarbon oil 700 (Sigma) and a membrane to prevent desiccation. Time-lapse imaging was performed on a Leica SP5 II confocal microscope at 21 °C, with 20 planes (z-step size: 0.45 µm) captured every 2 minutes for approximately 1 hour using a 63x objective (512 × 256 resolution). Maximum Z-projections were generated in Fiji (Schindelin et al., 2012) to create unprocessed movies (**Movies S1 and S2**).

### Image Processing

Nuclei were detected by Gaussian blurring, thresholding and applying a watershed algorithm. Transcription sites were detected as local maxima in 3D after applying a Difference-of-Gaussian filter. Transcription sites were assigned to the closest nucleus (**Supplementary File 1**, Program 1). Graphical representation of embryos and AP plots of embryos were generated in Matlab by identifying and quantifying nuclear puncta by extracting nuclei and associating puncta intensities with their respective nuclei. Fluorescence along the DV axis was summed and normalized to generate AP distribution plots (**Supplementary File 1**, Program 2). For transcription landscape profiling, data of several embryos are plotted on a space axis using Matlab (**Supplementary File 1**, Program 3).

For bursting analysis of live imaging movies, fluorescence time traces from individual nuclei were generated manually in Fiji. Nuclei traces and burst analysis was performed with Matlab. For each nucleus, bursts were identified as periods when the signal exceeds 5% of the maximum trace intensity. Mean burst amplitude is determined by averaging the maximum signal intensities within each burst. Mean intensity corresponds to the average signal across the entire trace (**Supplementary File 1**, Program 4).

## Movie Availability and Legends

Movies can be accessed through the following link: https://owncloud.gwdg.de/index.php/s/6eJOI2JvyoYNffU

**Movie S1. MS2-MCP live imaging of 3&7&2 enhancer reporter line.** Live imaging of a “3&7&2>MS2-*yellow*; MCP-GFP; Histone-RFP” embryo from early NC14 to late NC14 at stripe position 2. Histone-RFP marks nuclei; MS2 loops recruit MCP-GFP to active transcription sites, producing strong GFP puncta at stripe 2, reflecting enhancer-driven mRNA transcription. Anterior is left.

**Movie S2. MS2-MCP live imaging of 3&7-su(Hw) upstream-2 enhancer reporter line.** Live imaging of a “3&7-su(Hw) upstream-2>MS2-*yellow*; MCP-GFP; Histone-RFP” embryo from early NC14 to late NC14 at stripe position 2. Histone-RFP marks nuclei; MS2 loops recruit MCP-GFP to active transcription sites, producing slightly weaker GFP puncta at stripe 2 compared to the “3&7&2>MS2-*yellow*; MCP-GFP; Histone-RFP” embryo. Burst frequency and intensity is significantly different (see **Figure 6C** for details). Anterior is left.

**Supplementary Table 1.**
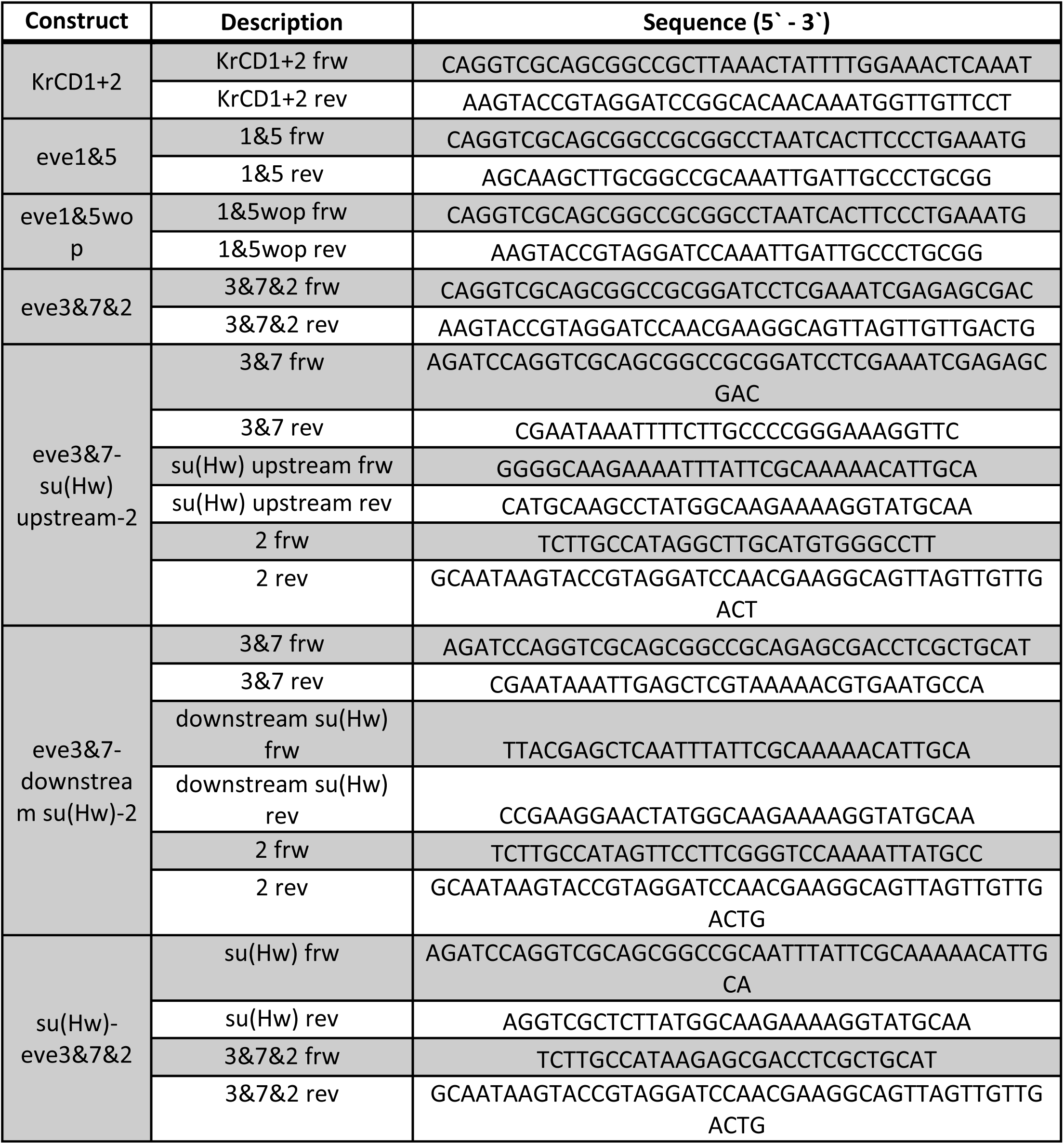
List of used primers. Frw: forward, rev: reverse.

**Supplementary Table 2.** LOT numbers for HCR probe sets. Landscape profiling: individual probes of a probe set separately delivered allowing for eRNA transcription detection with a resolution down to 500 bp.

| LOT Number | Target | HCR Amplifier |
| --- | --- | --- |
| PRA987 | *yellow* mRNA | B1 |
| PRB336 | *Kr* mRNA | B1 |
| PRB337 | KrCD1 sense eRNA | B2 |
| PRB338 | KrCD1 anti-sense eRNA | B2 |
| PRB339 | KrCD2 sense eRNA | B3 |
| PRB340 | KrCD2 anti-sense eRNA | B3 |
| PRB341 | *eve* mRNA | B1 |
| PRB874 | *eve* stripe3&7 sense eRNA -<br>landscape profiling | B2 |
| PRB875 | *eve* intergenic region between<br>*eve* stripe 3&7 and stripe 2<br>sense eRNA - landscape<br>profiling | B2 |
| PRB876 | *eve* stripe2 sense eRNA -<br>landscape profiling | B2 |
| PRB877 | *eve* intergenic region between<br>*eve* stripe 2 and *eve*<br>transcription start site sense<br>eRNA - landscape profiling | B2 |
| PRB883 | *eve* stripe1 sense eRNA | B2 |
| PRB884 | *eve* stripe5 sense eRNA | B2 |
| PRC021 | *gt* mRNA | B3 |
| PRC022 | gt(-1) sense eRNA | B1 |
| PRC023 | gt(-1) anti-sense eRNA | B1 |
| PRC024 | gt(-3) sense eRNA | B2 |
| PRC025 | gt(-3) anti-sense eRNA | B2 |
| PRC654 | *yellow* mRNA | B2 |
| PRC655 | *yellow* mRNA | B3 |
| PRC668 | pBPhi exo sense eRNA | B3 |
| PRD263 | *Kr* locus sense eRNA -<br>landscape profiling | B2 |
| PRD266 | *Kr* locus anti-sense eRNA -<br>landscape profiling | B3 |
| PRE719 | pBPhi exo sense eRNA -<br>landscape profiling | B2 |

---

**Supplementary Figure 1.**
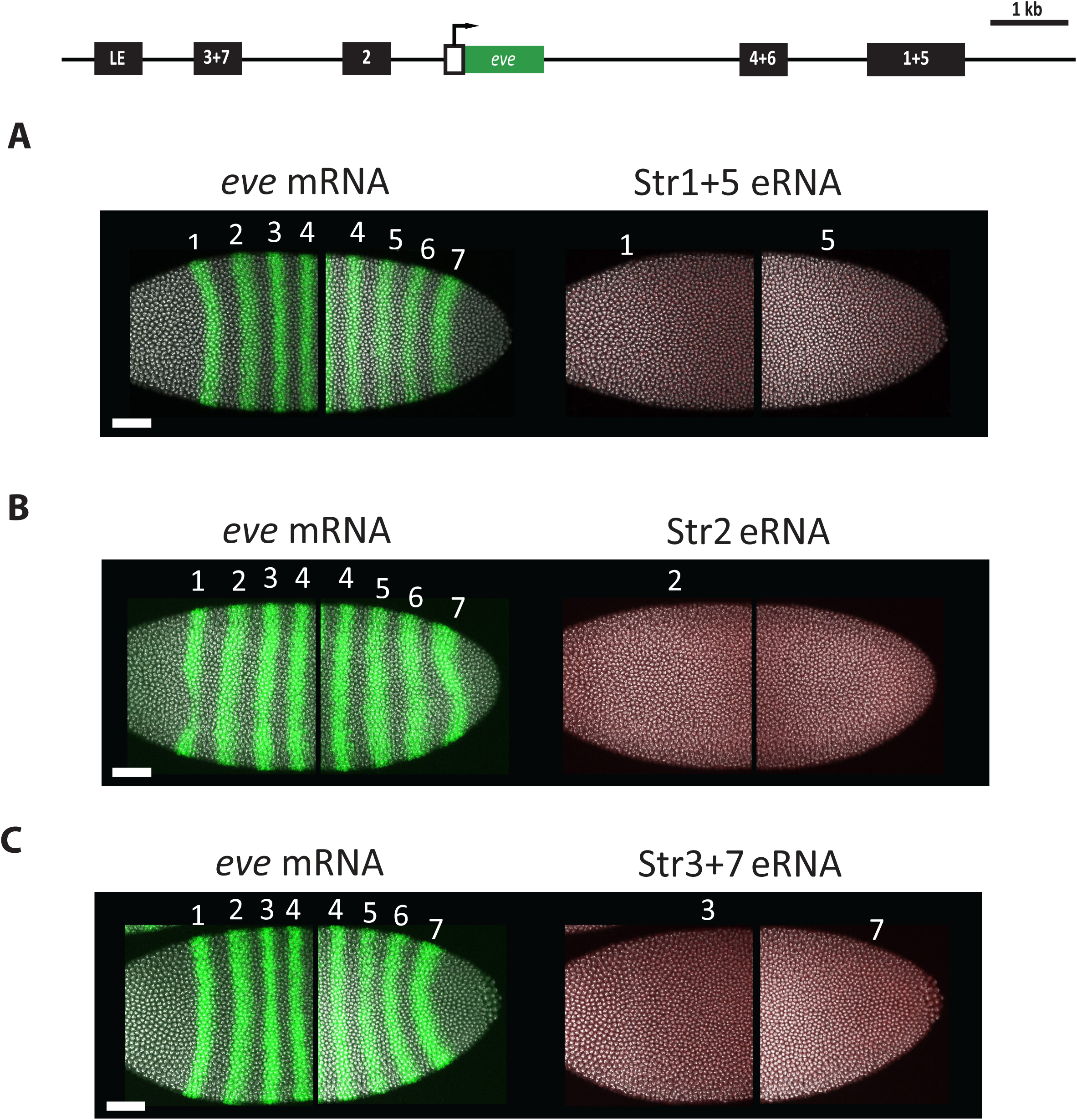
Spatial expression of eRNAs from *eve* enhancers at endogenous locus. mRNA (green) and eRNA (red) transcripts were visualized together in the endogenous *eve* locus in whole *Drosophila* embryos at nuclear cycle 14 (NC14) using *in situ* HCR staining. Nuclei are shown in grey. (**A**) *eve* 1+5 enhancer, (**B**) *eve* 2 enhancer, and (**C**) *eve* 3+7 enhancer showed no detectable eRNA transcripts. Anterior is left, dorsal is up. Scale bars: 50 µm.

## Supplementary File 1 – Fiji and Matlab code

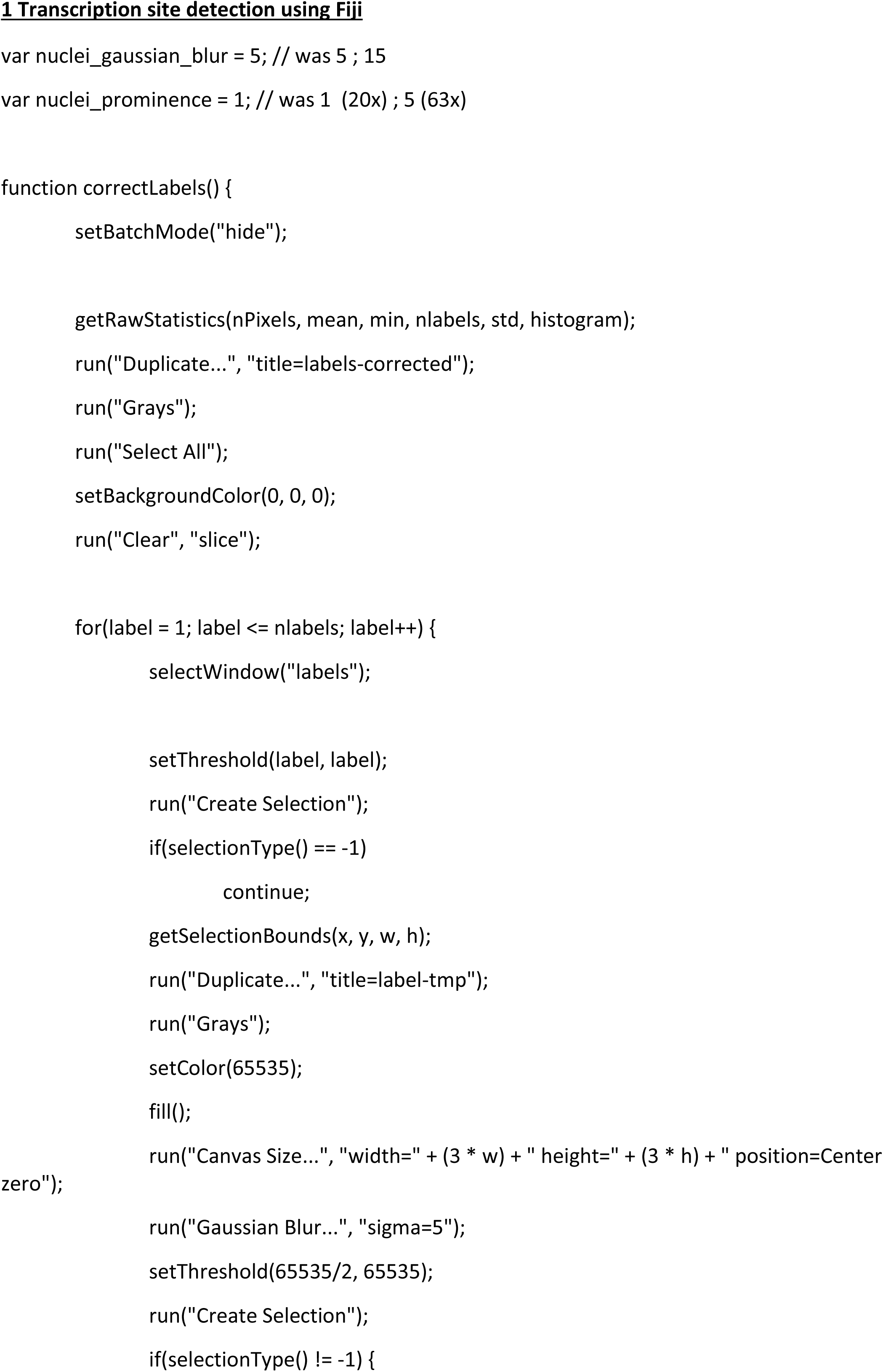
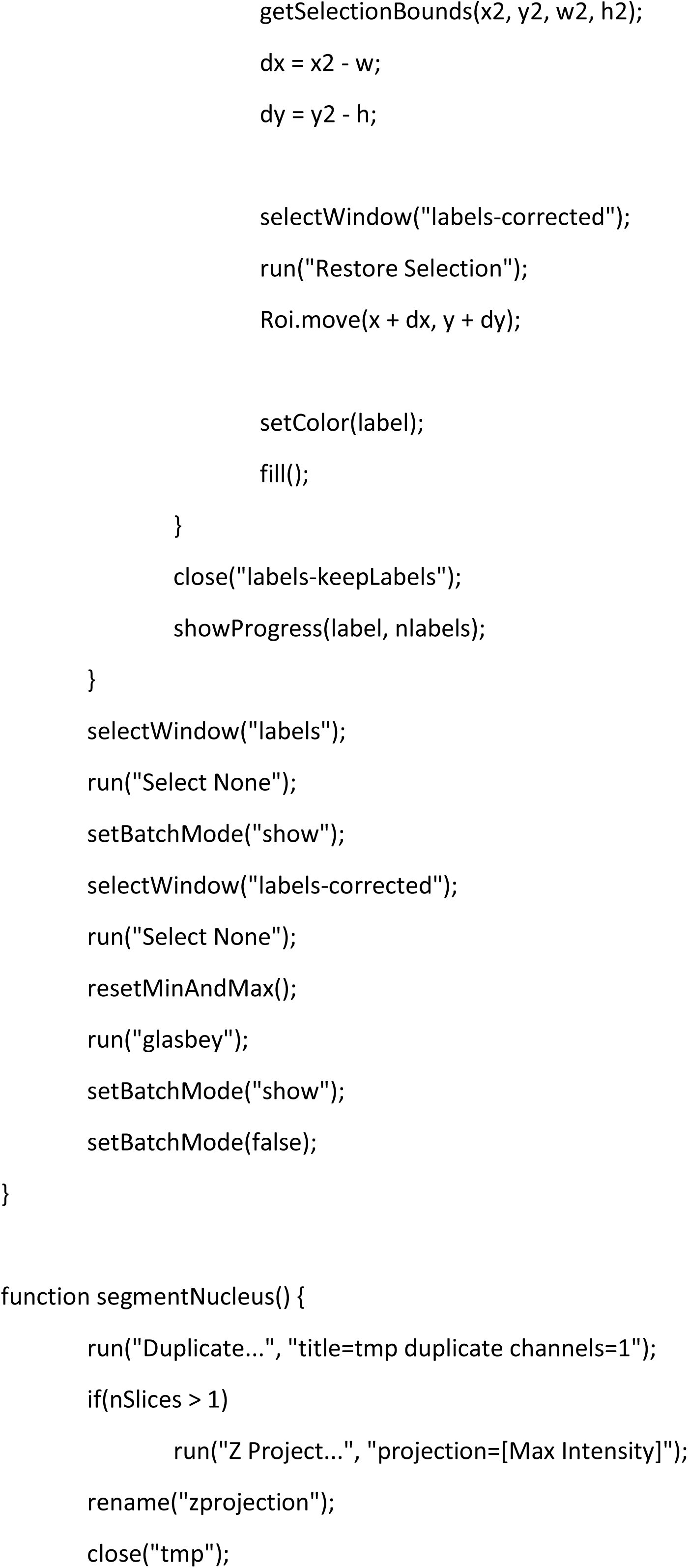
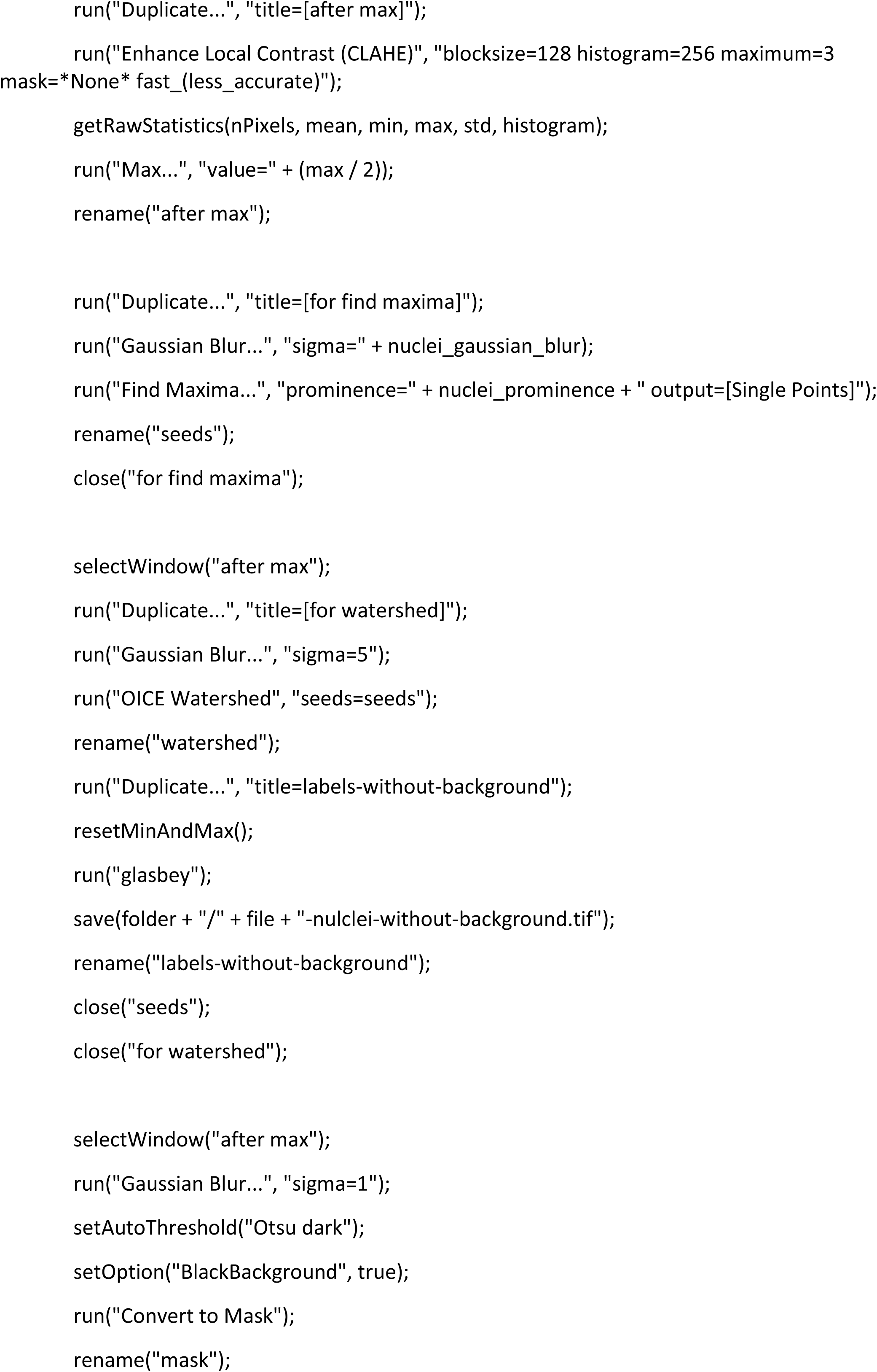
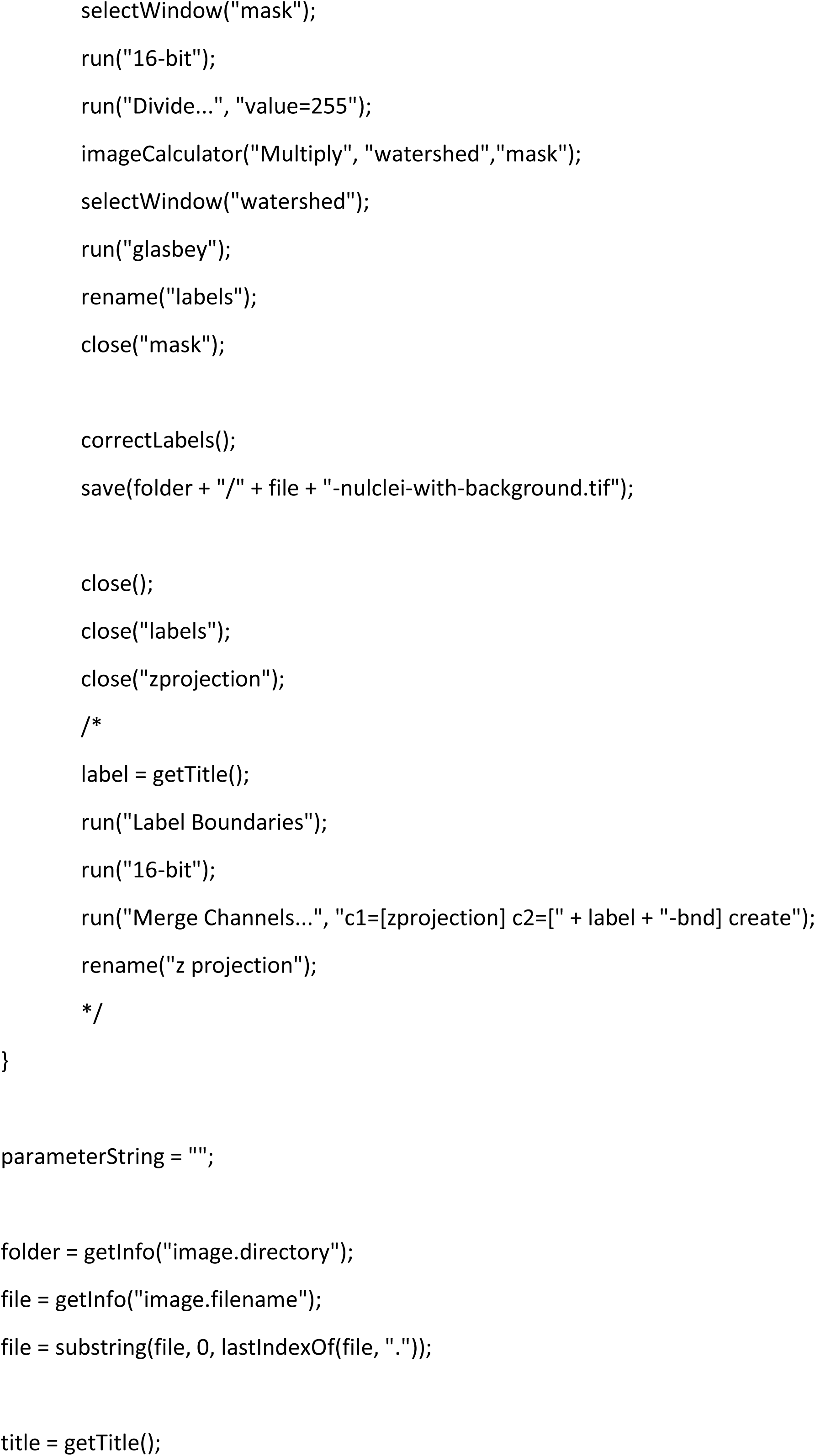
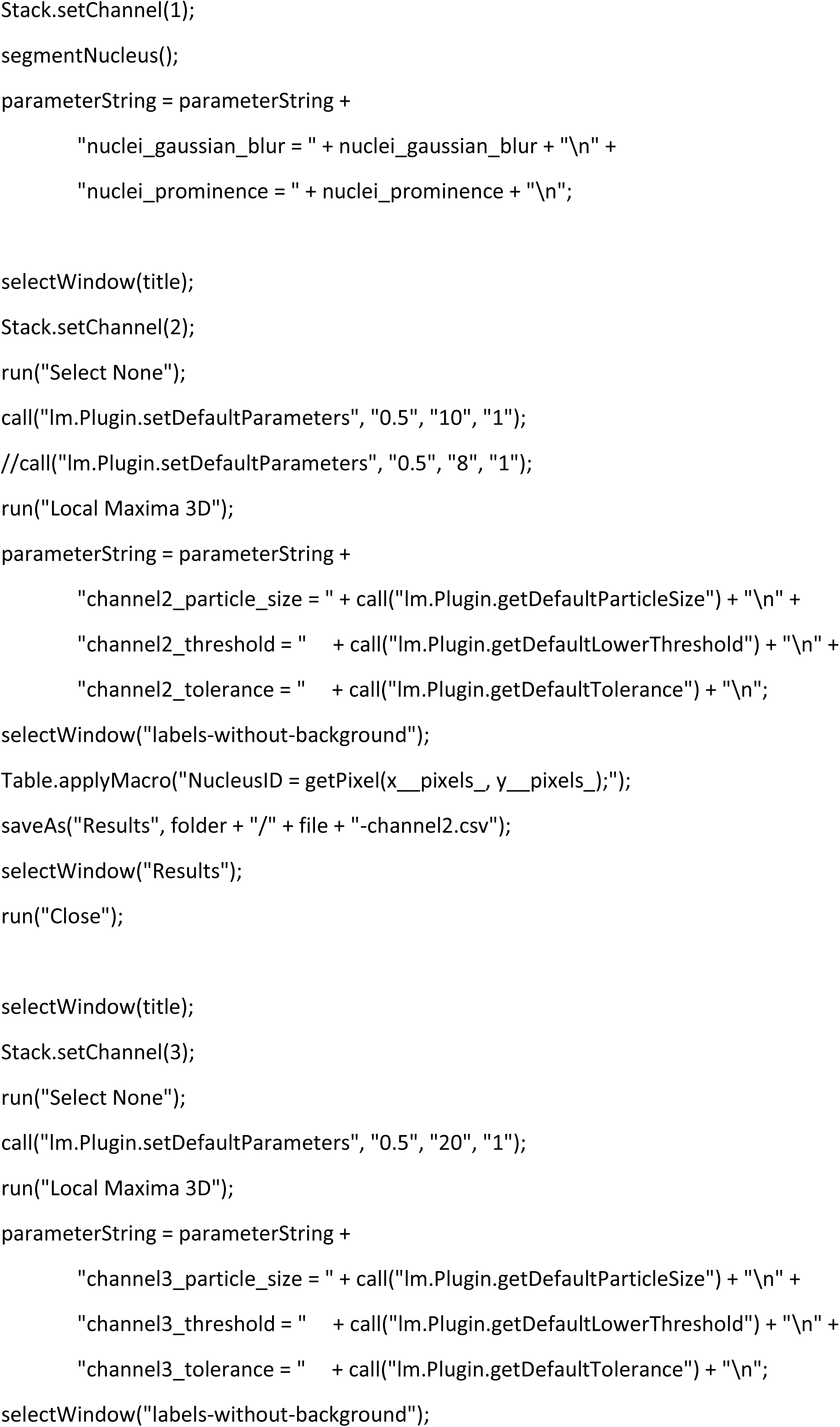
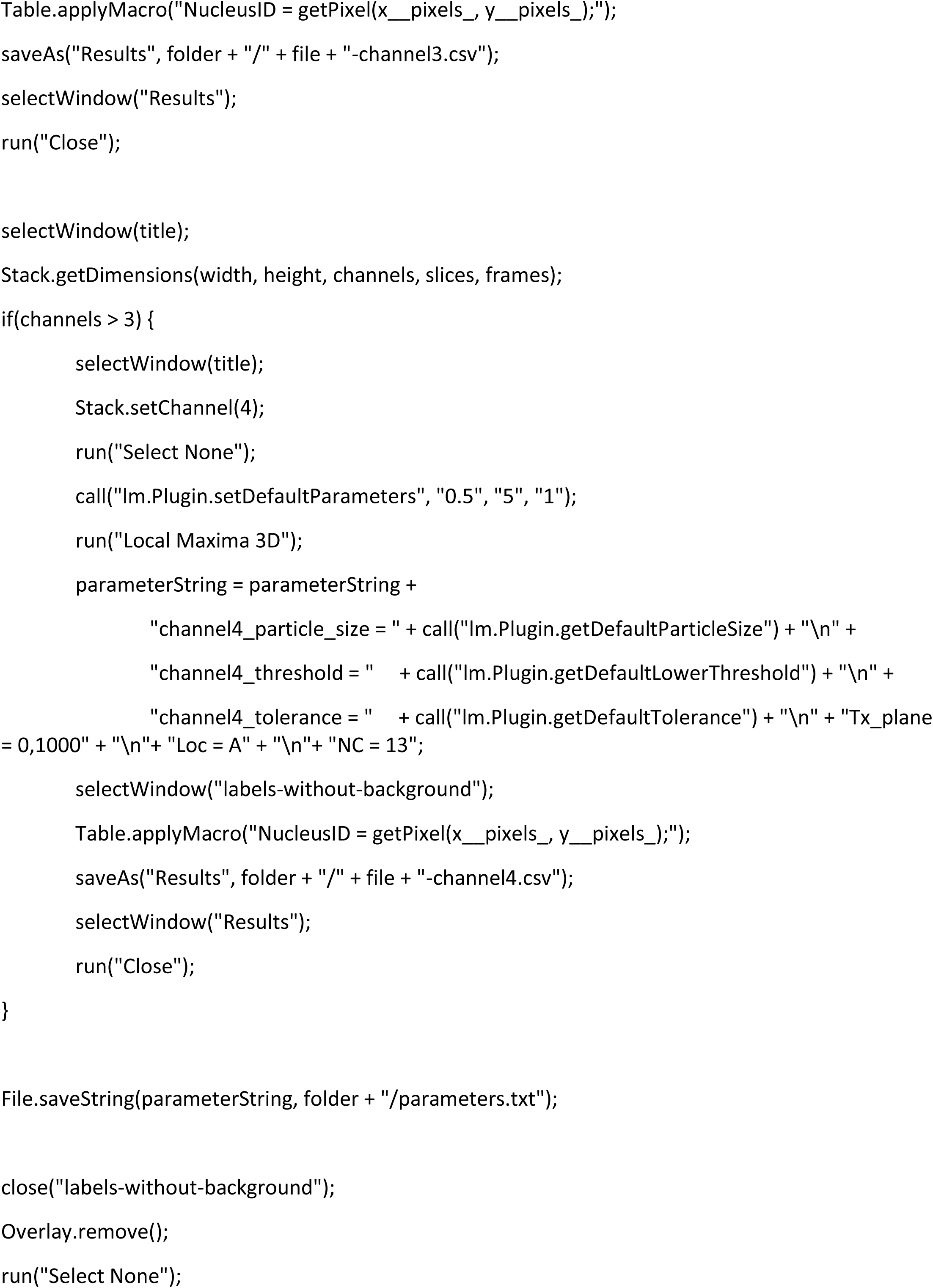
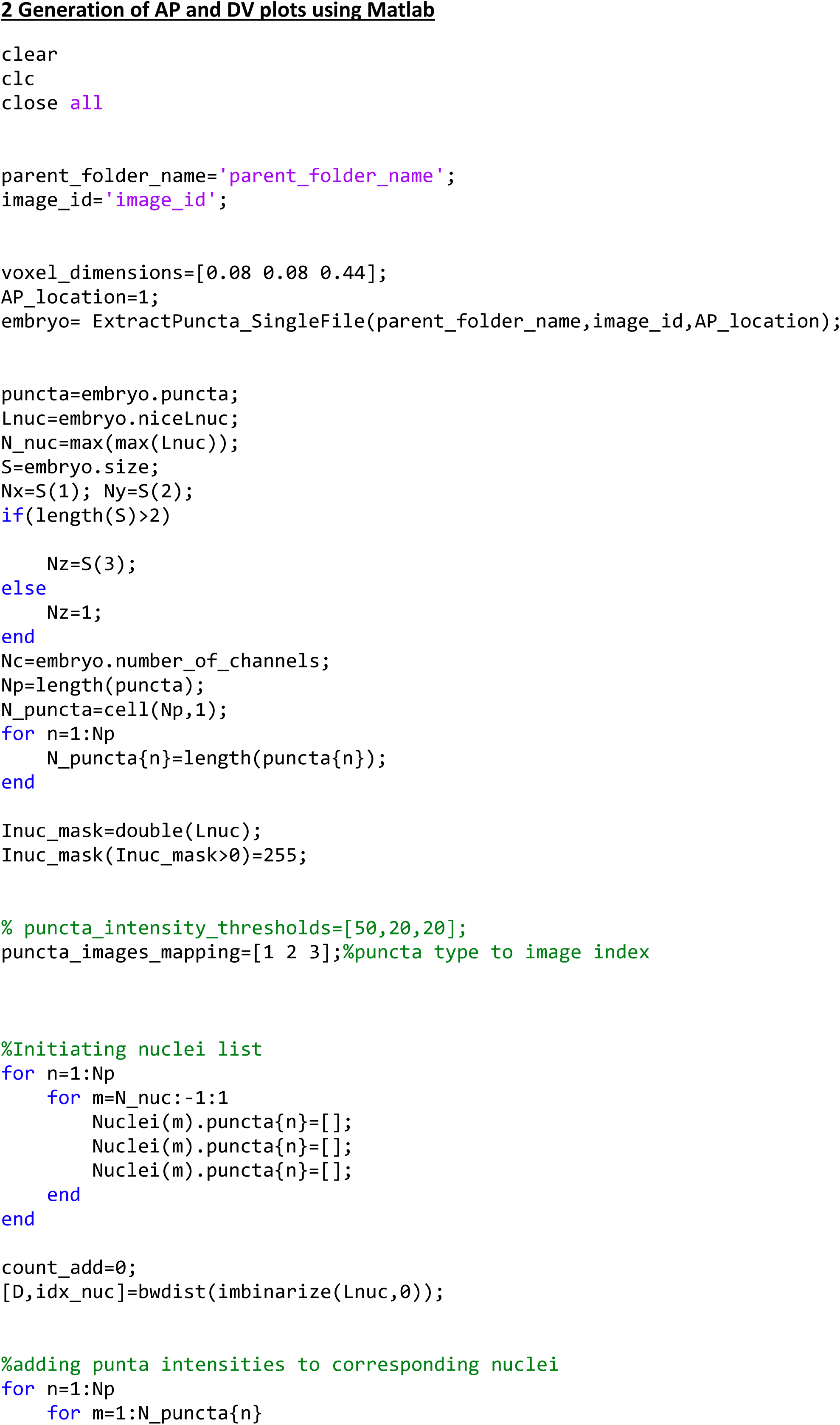
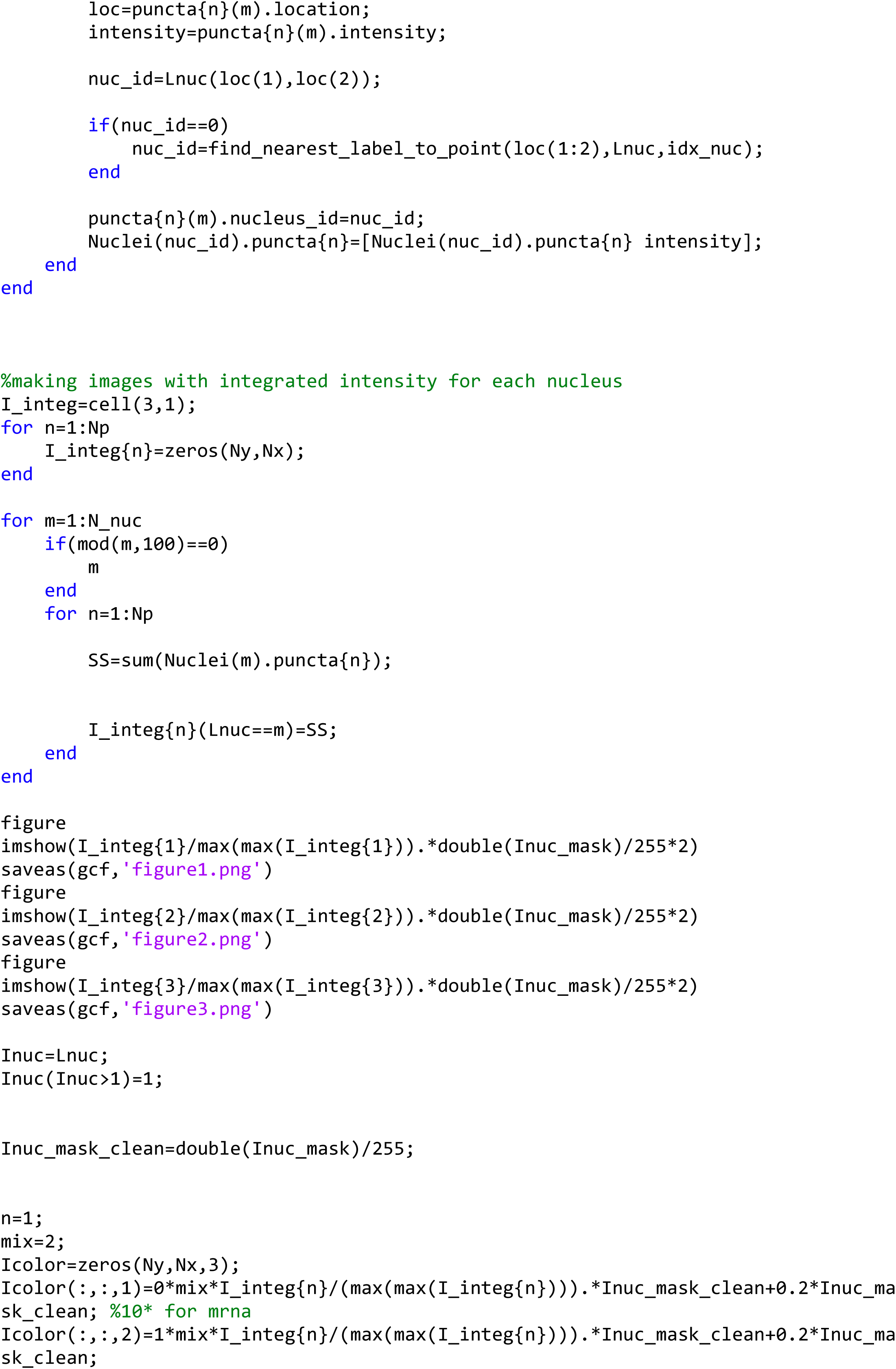
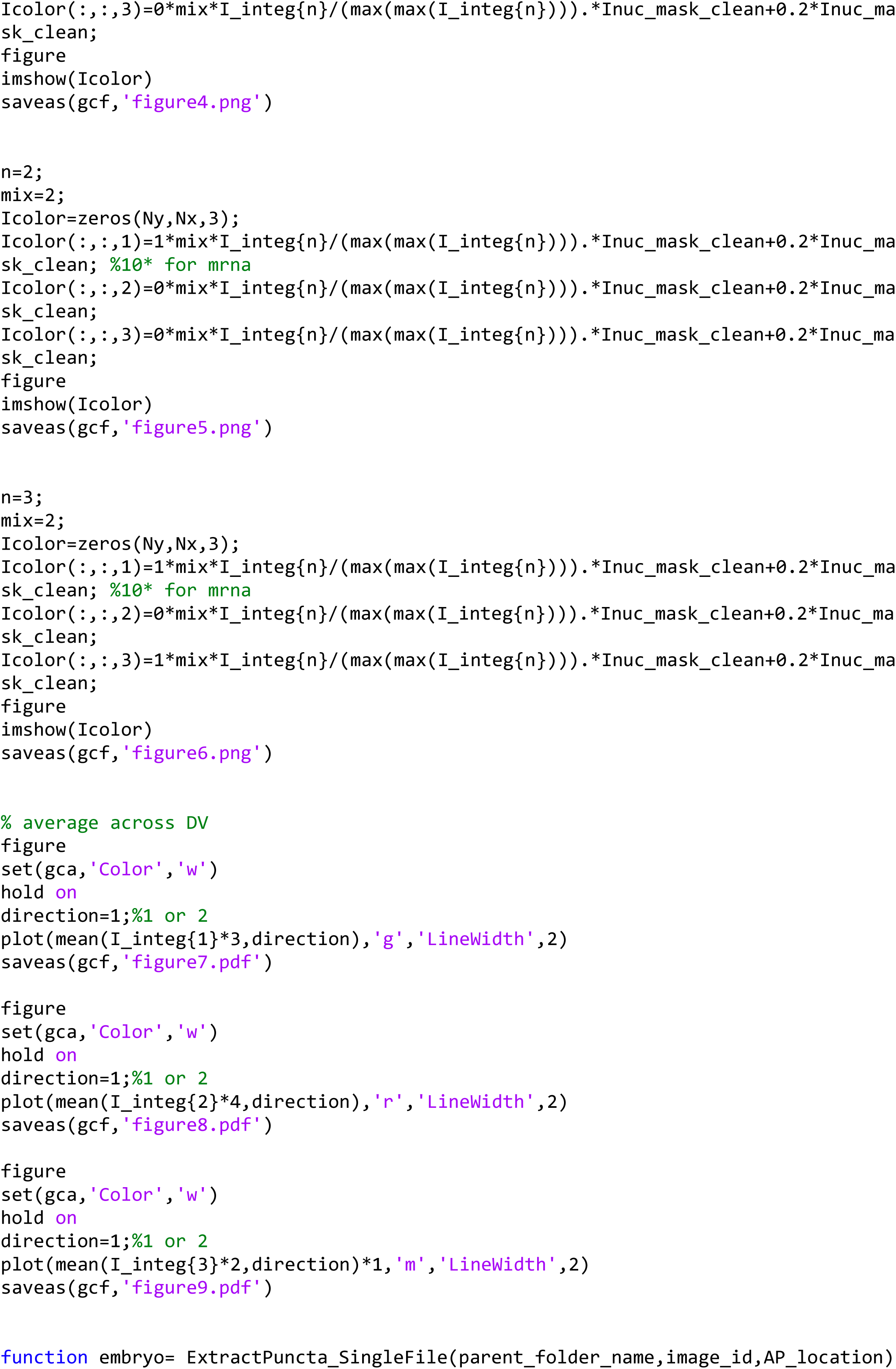
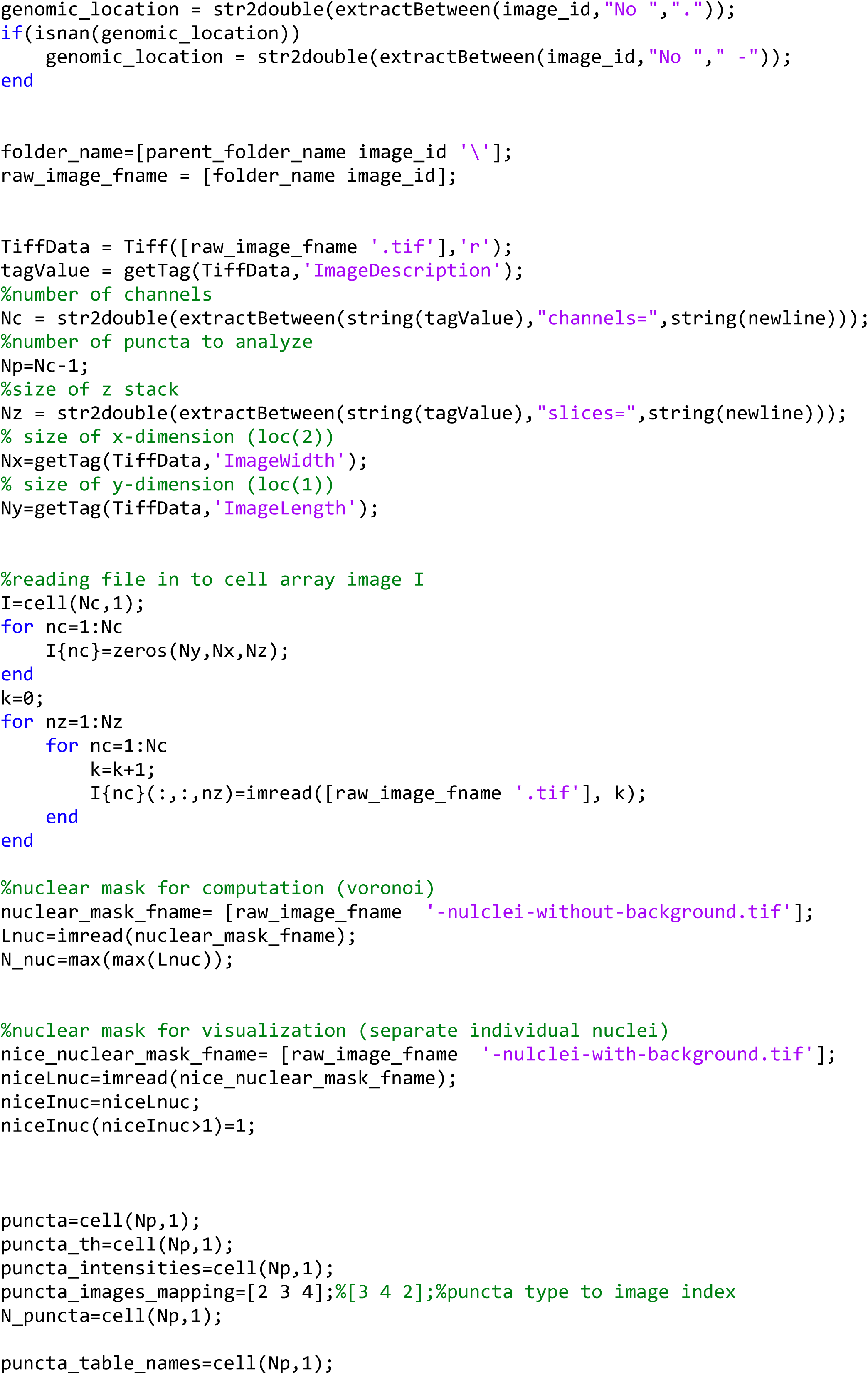
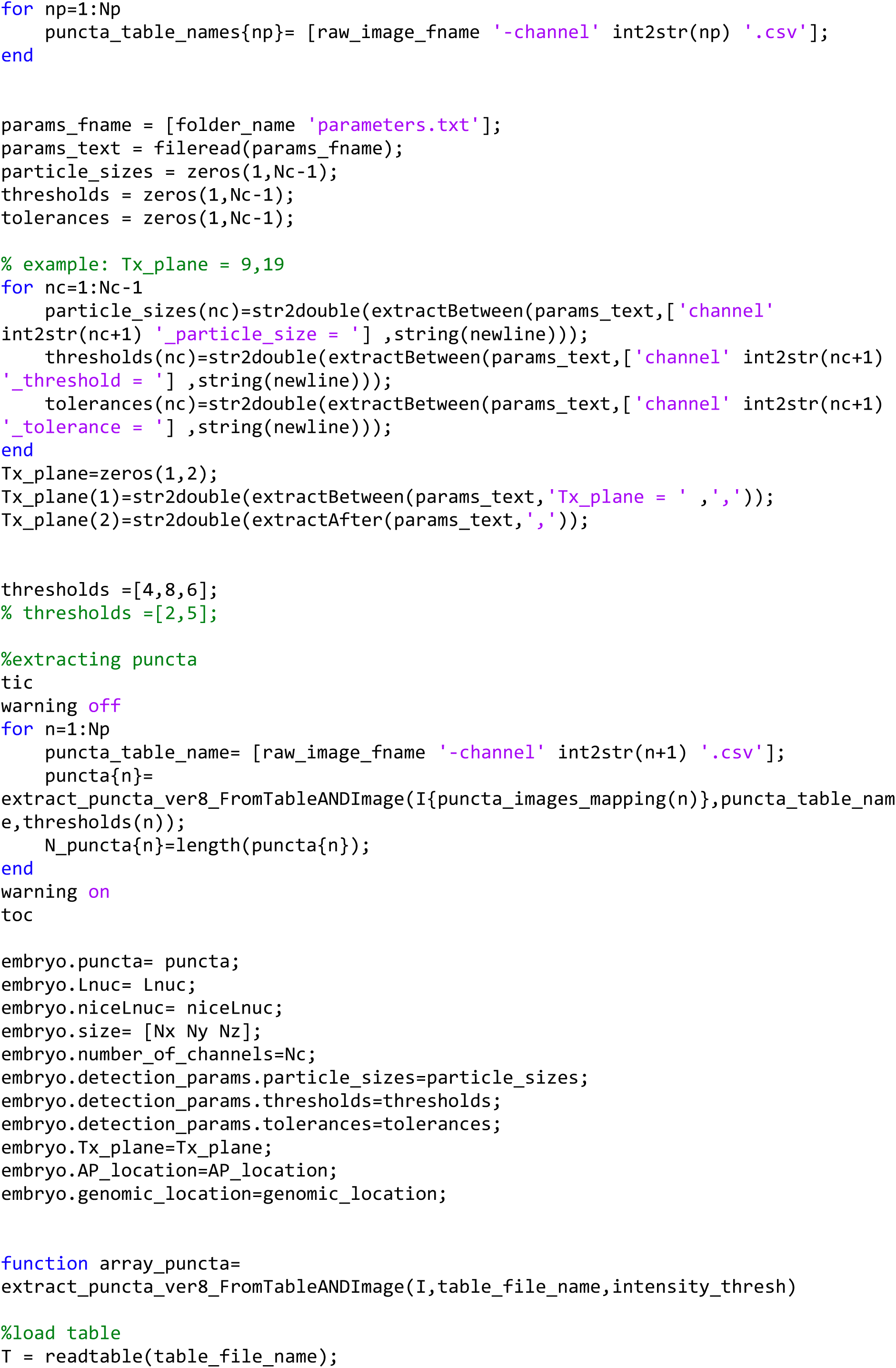
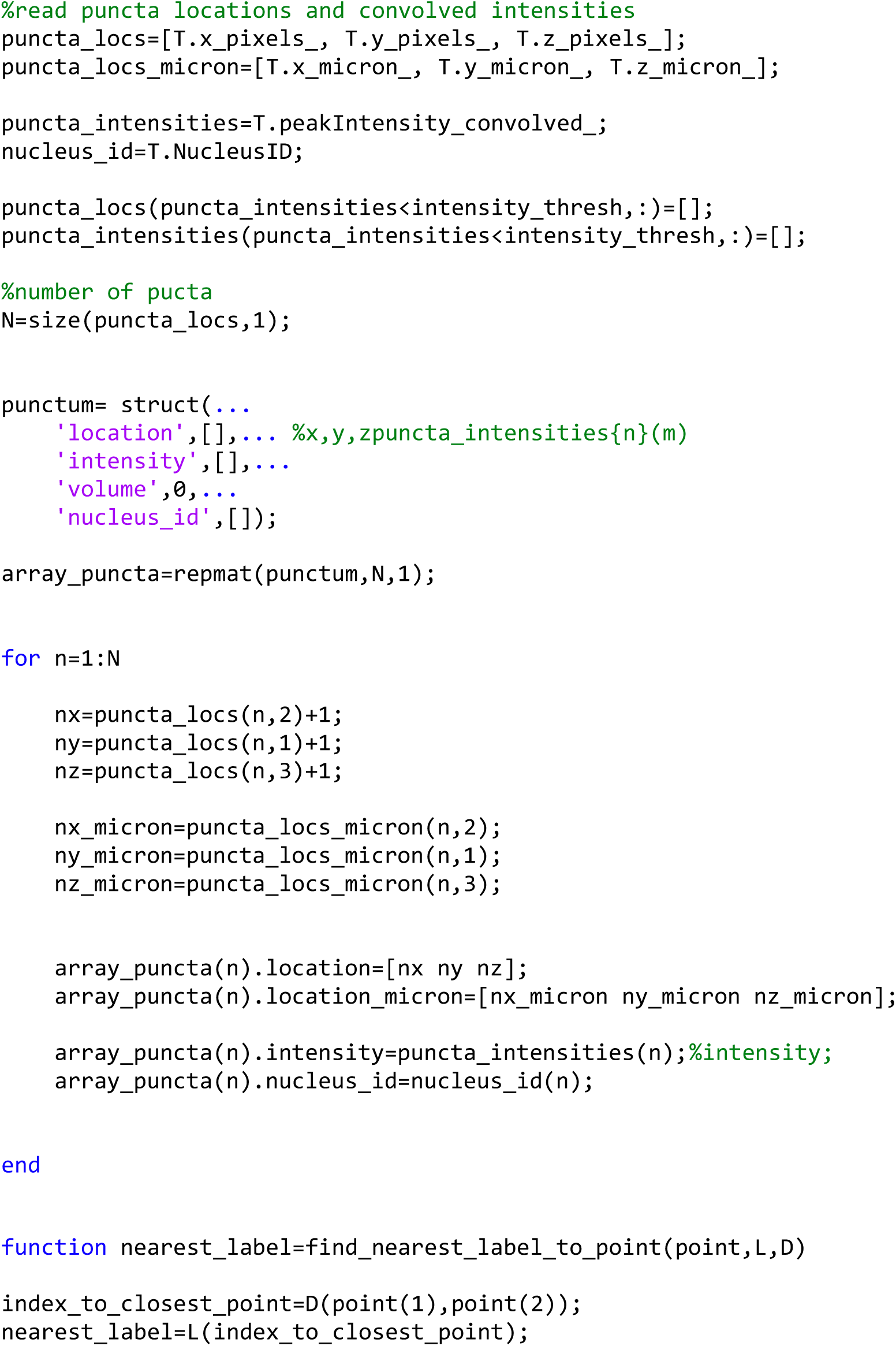
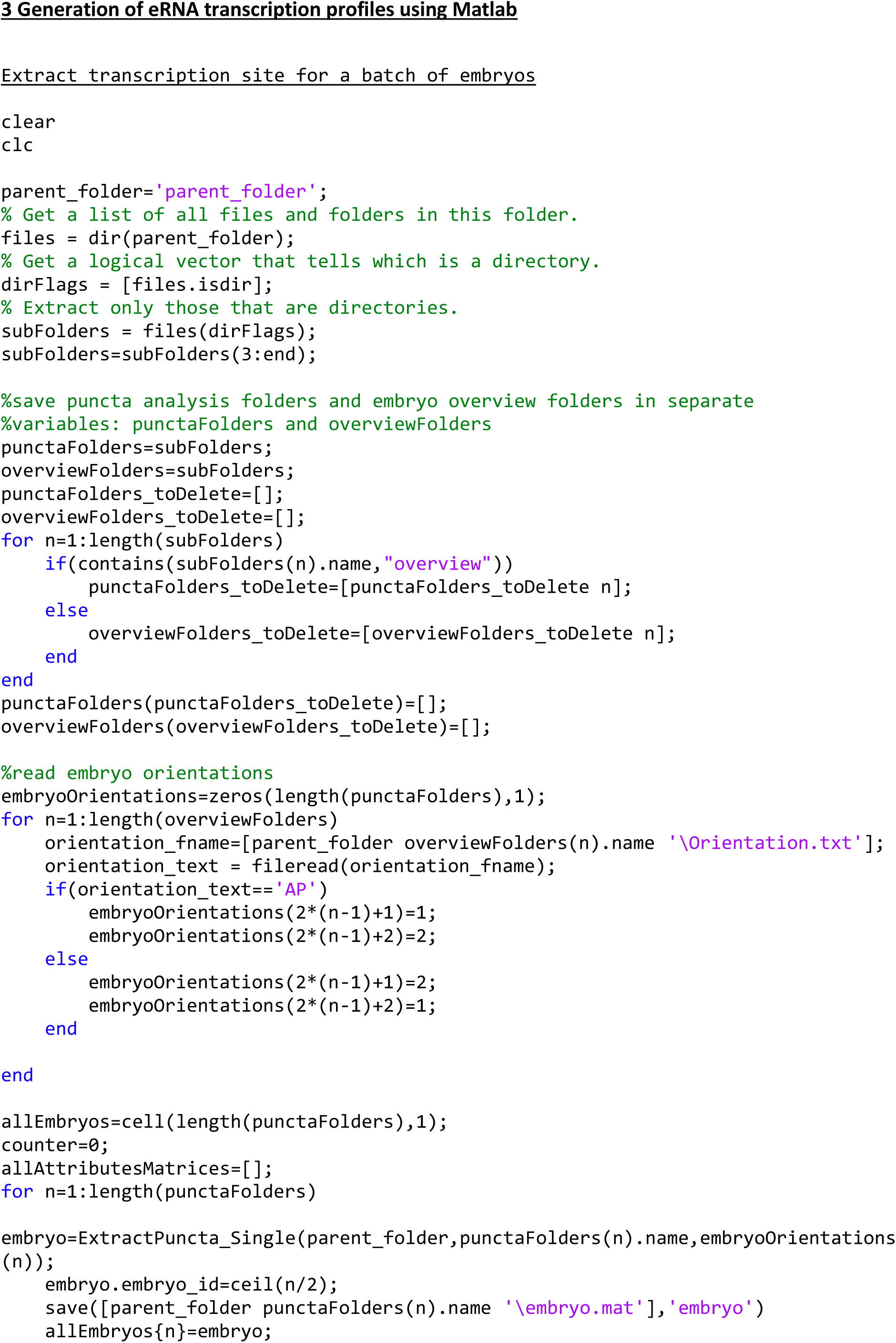
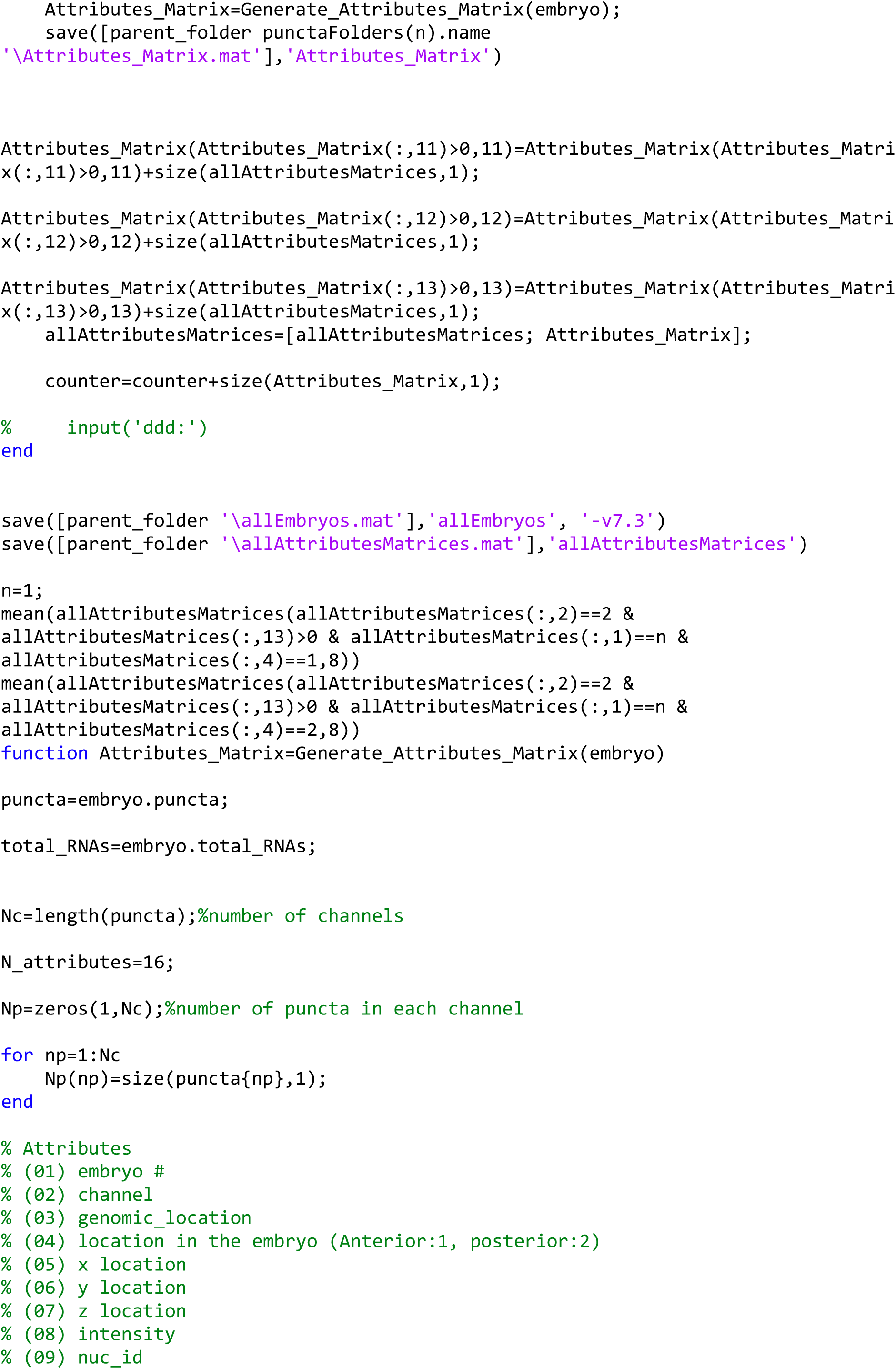
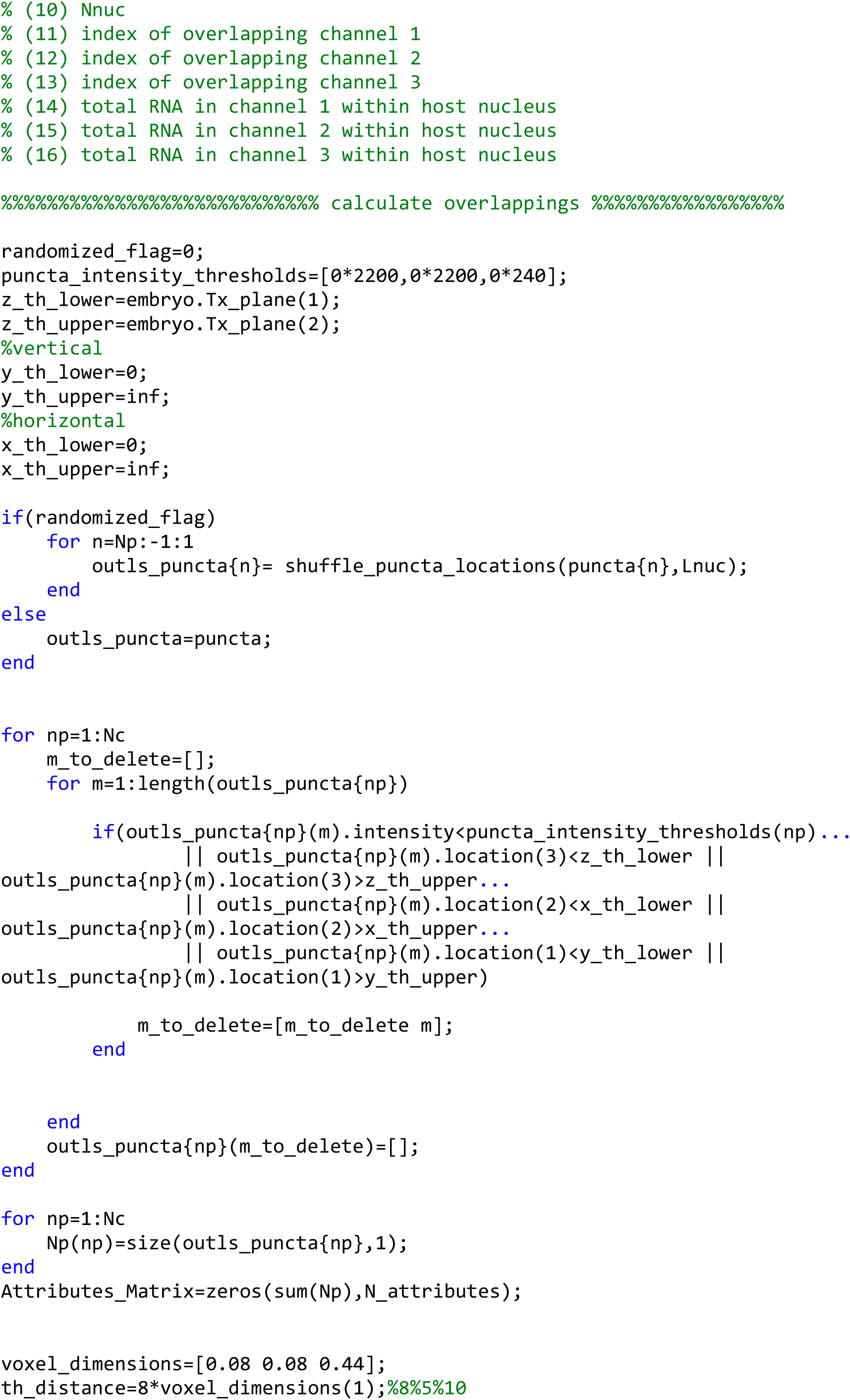
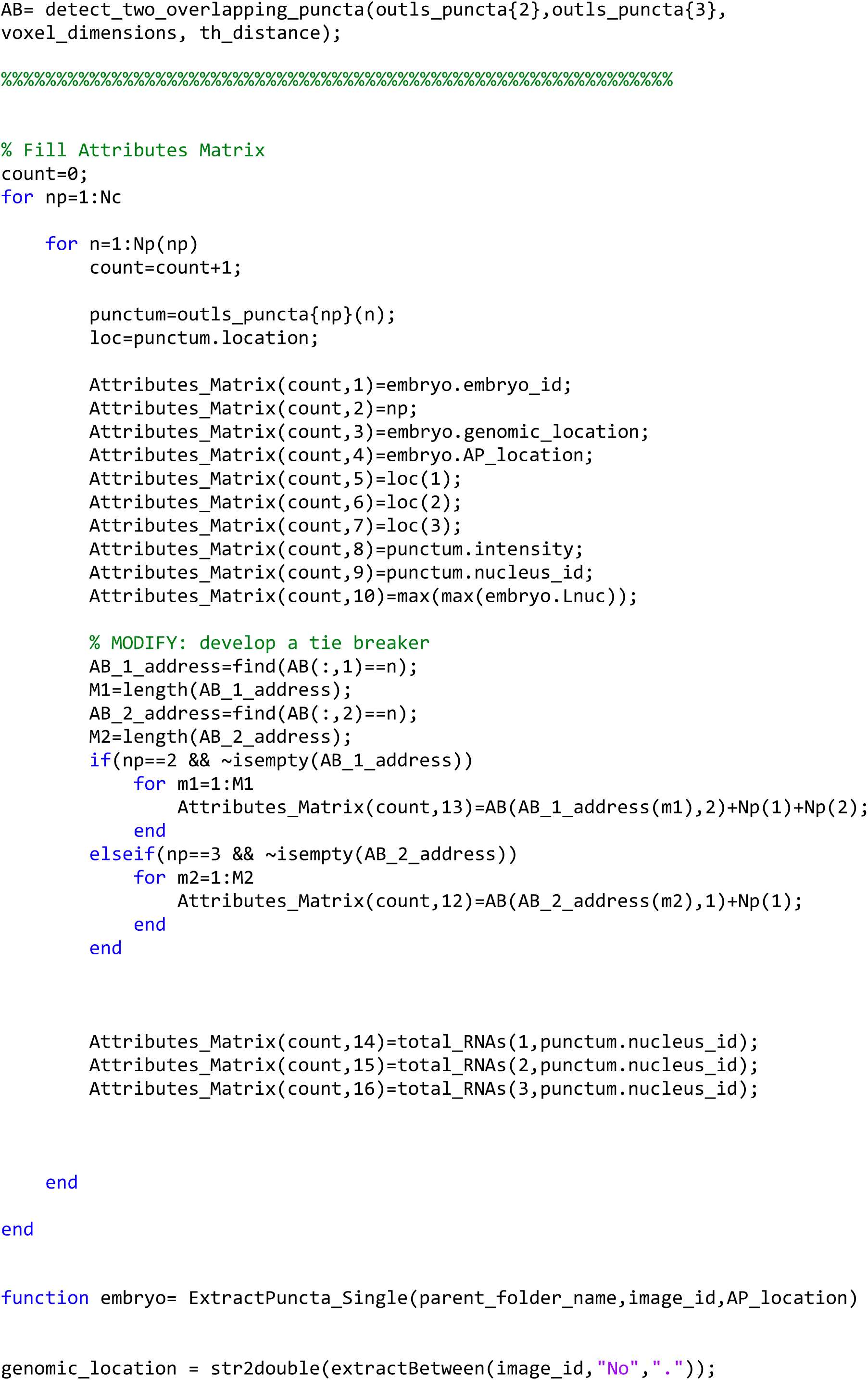
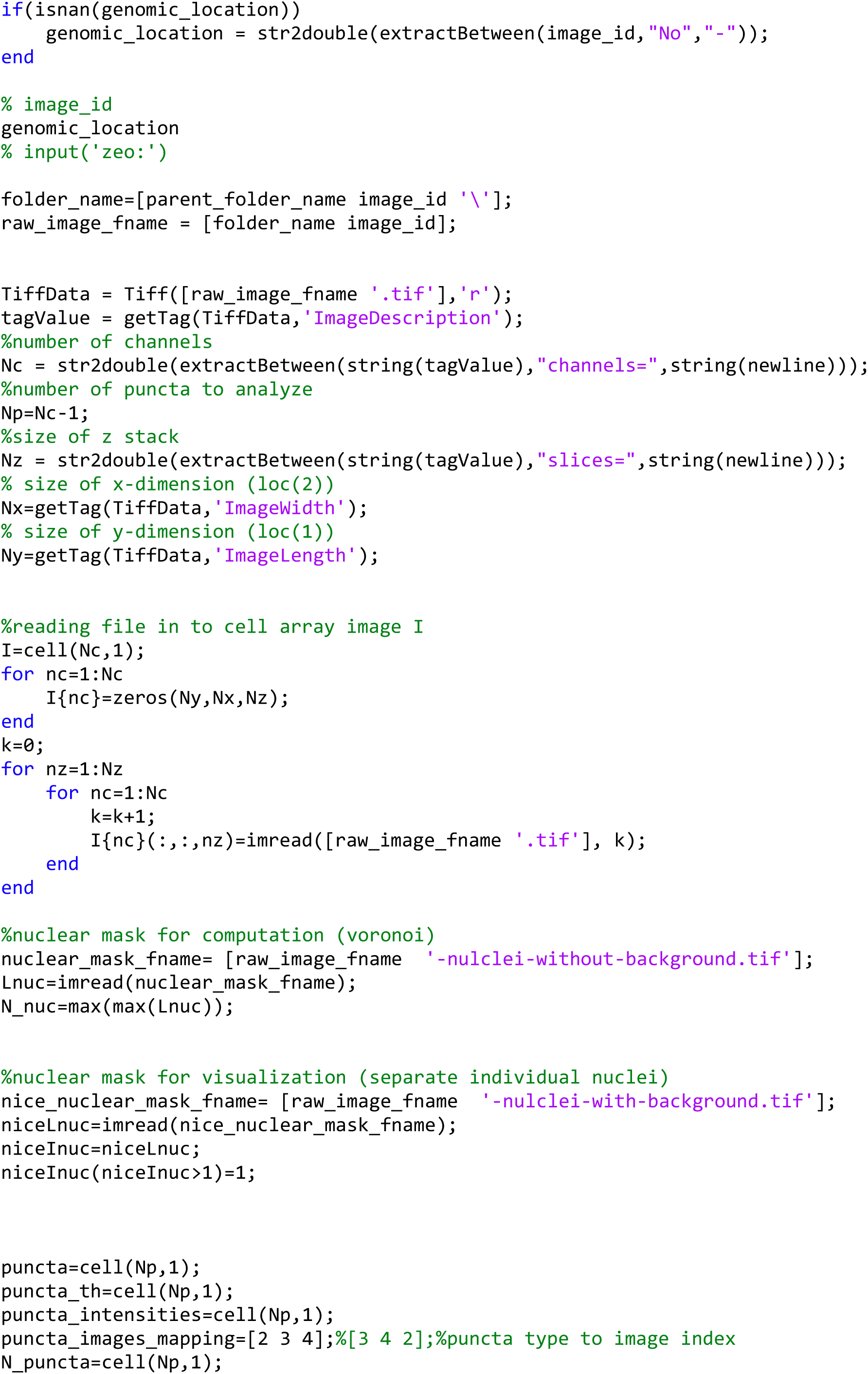
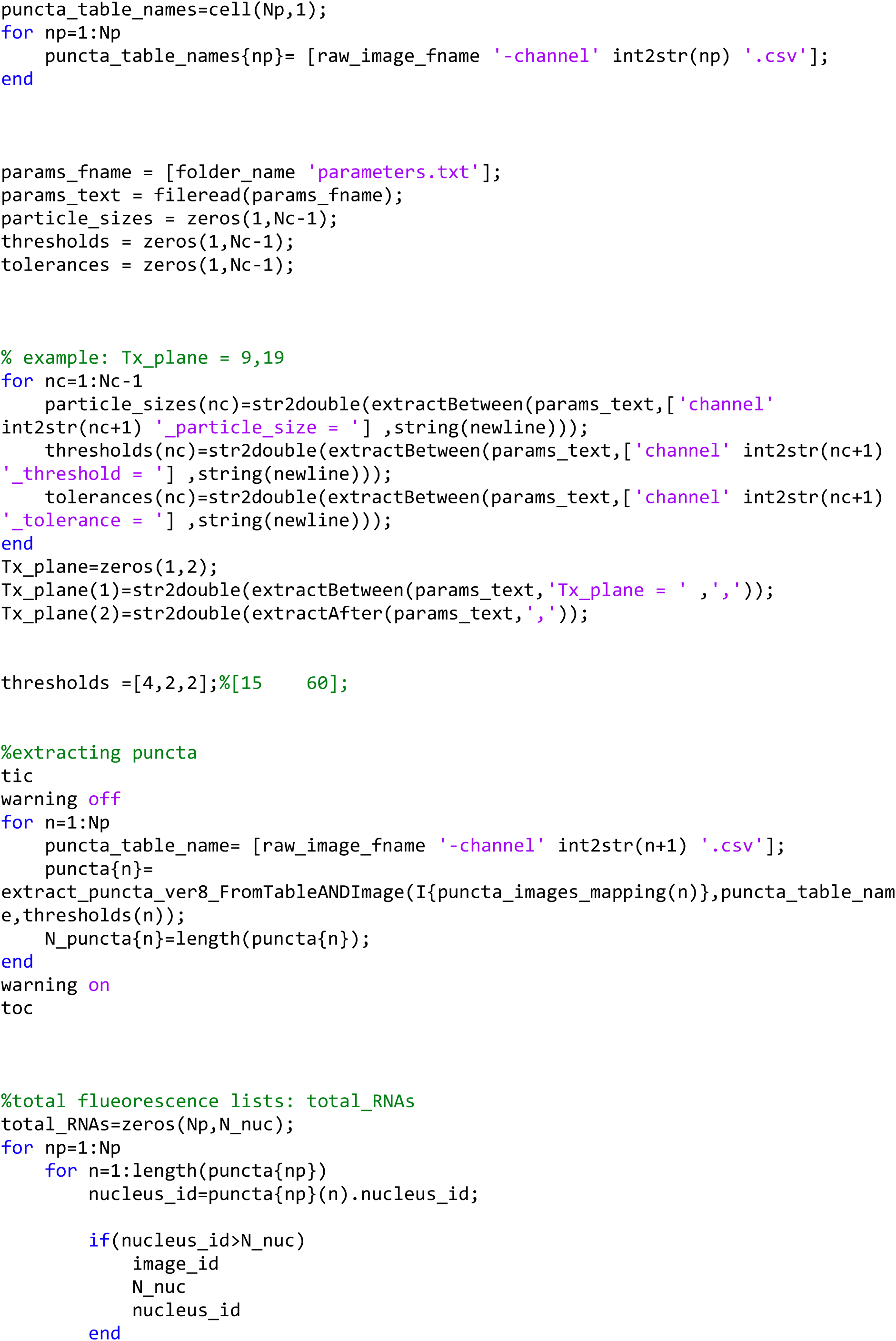
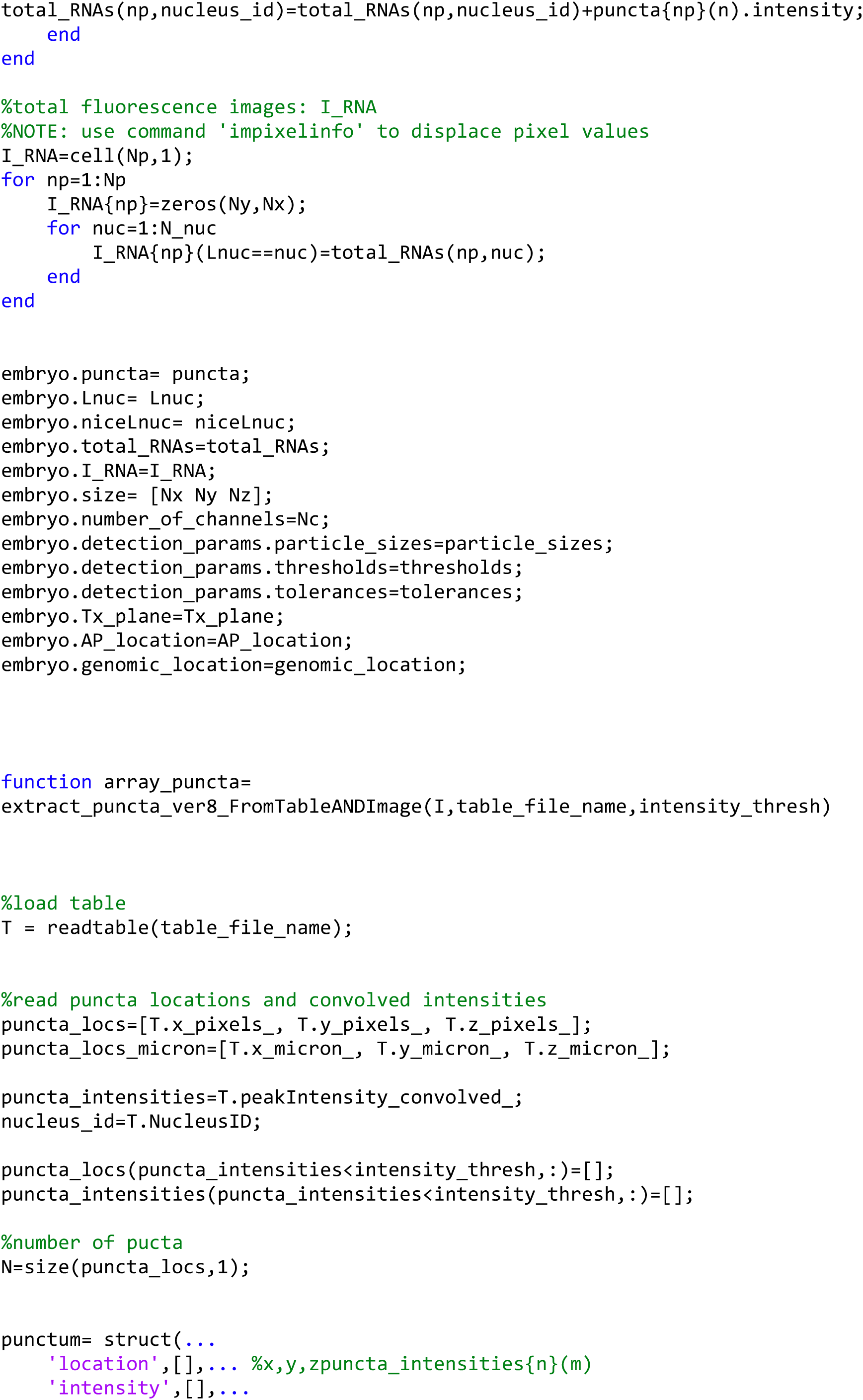
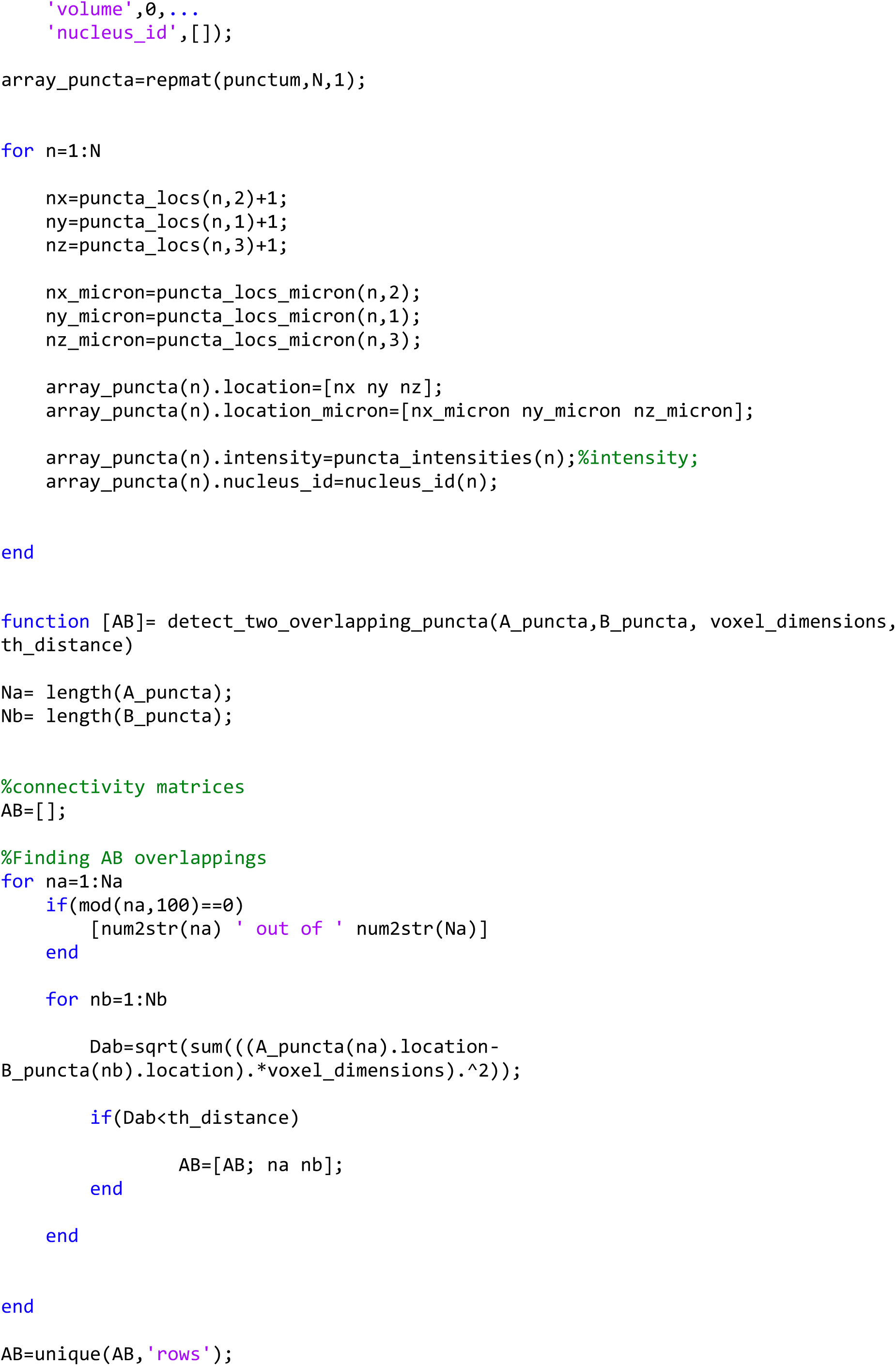
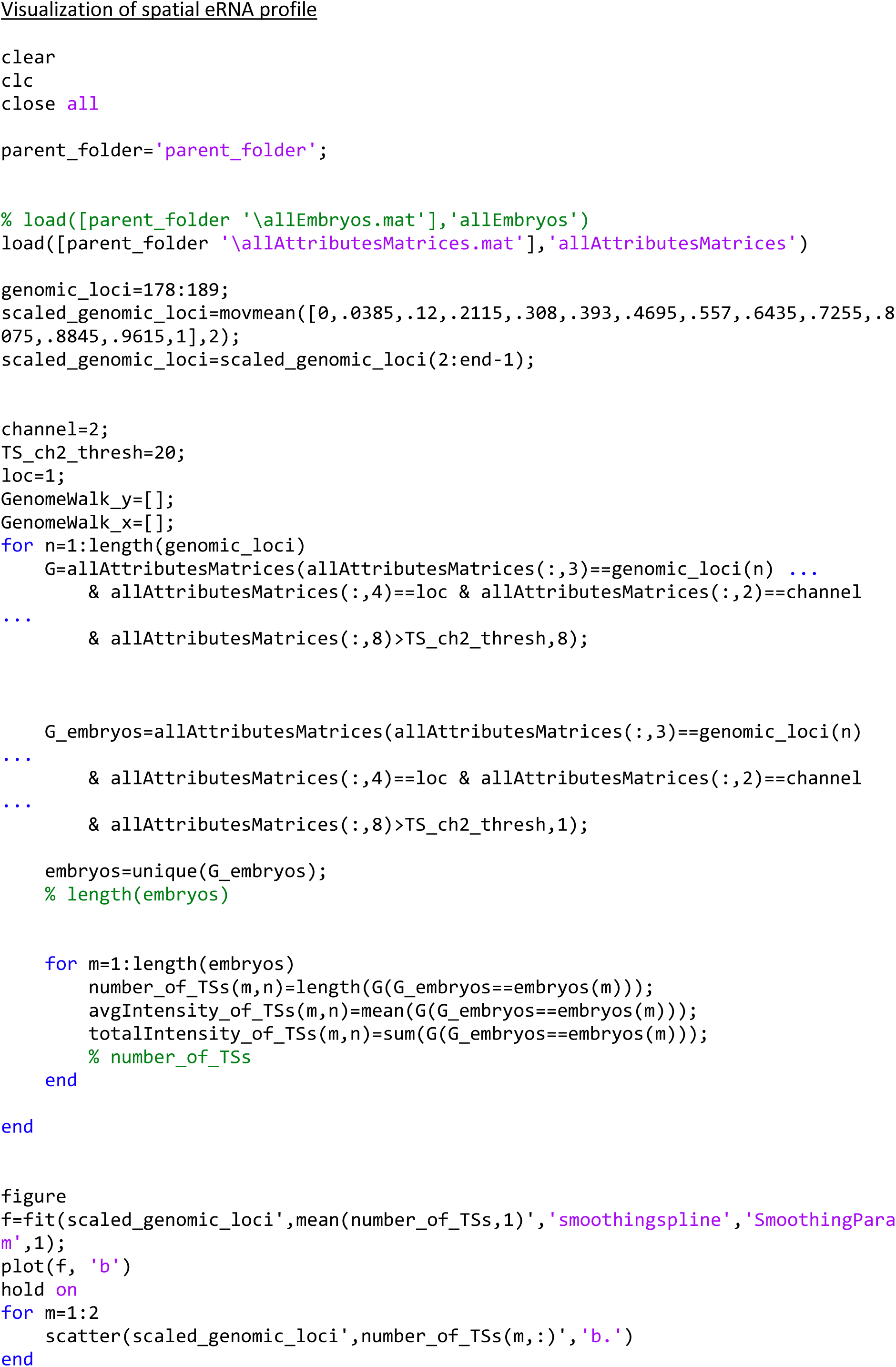
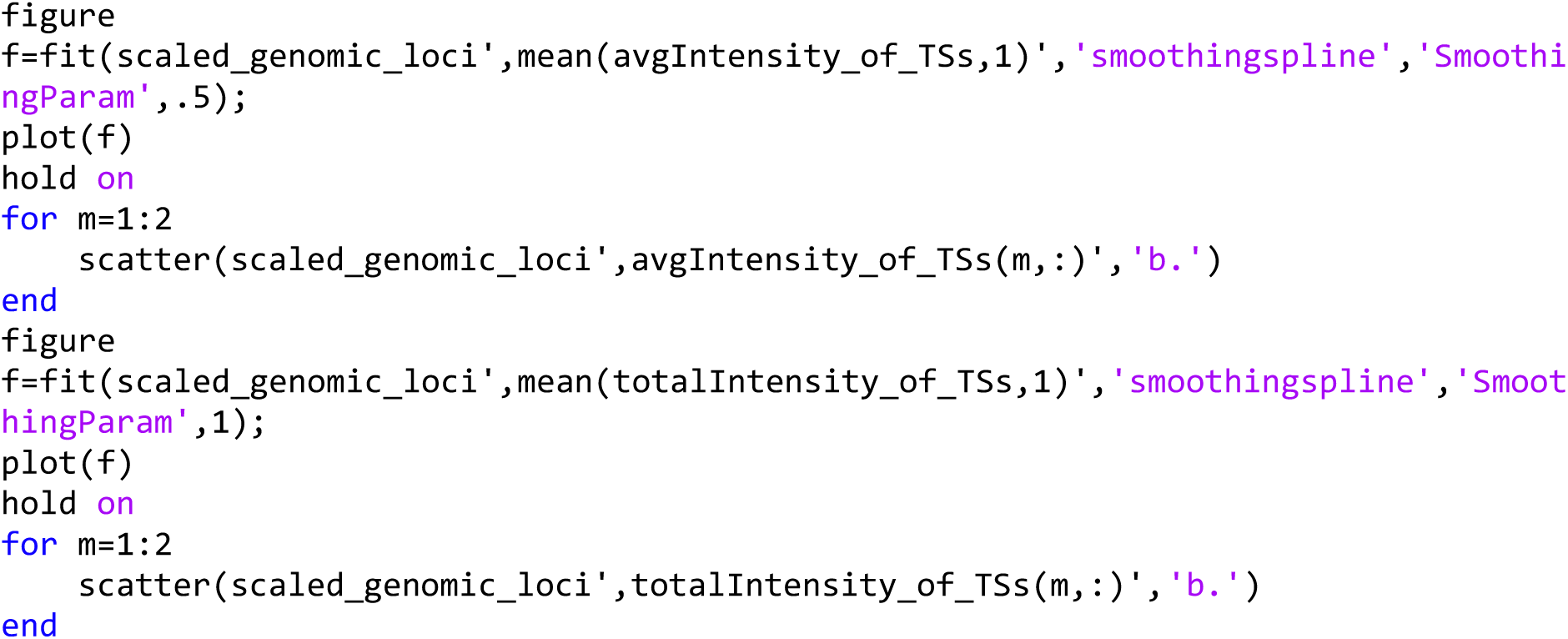
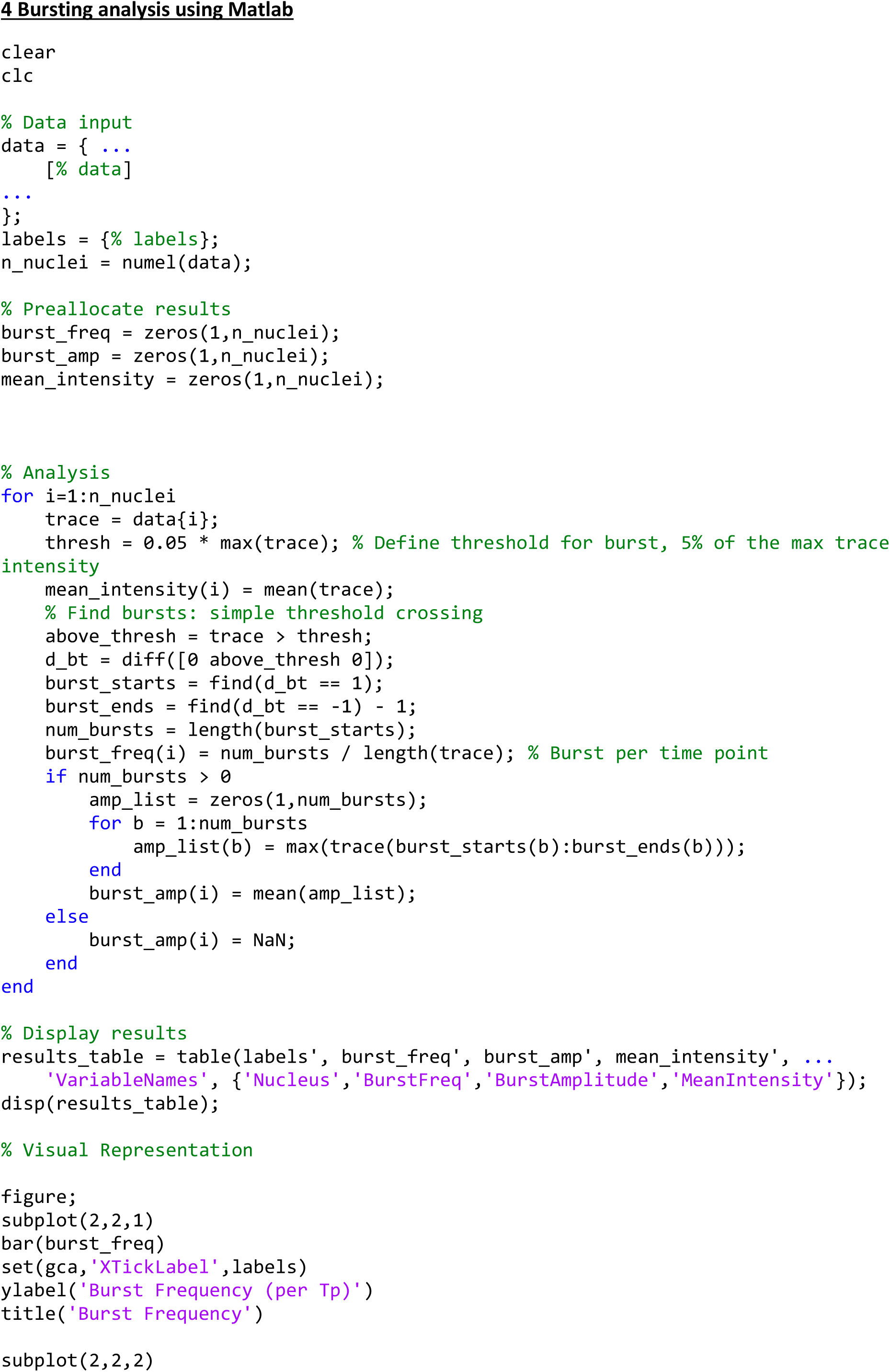
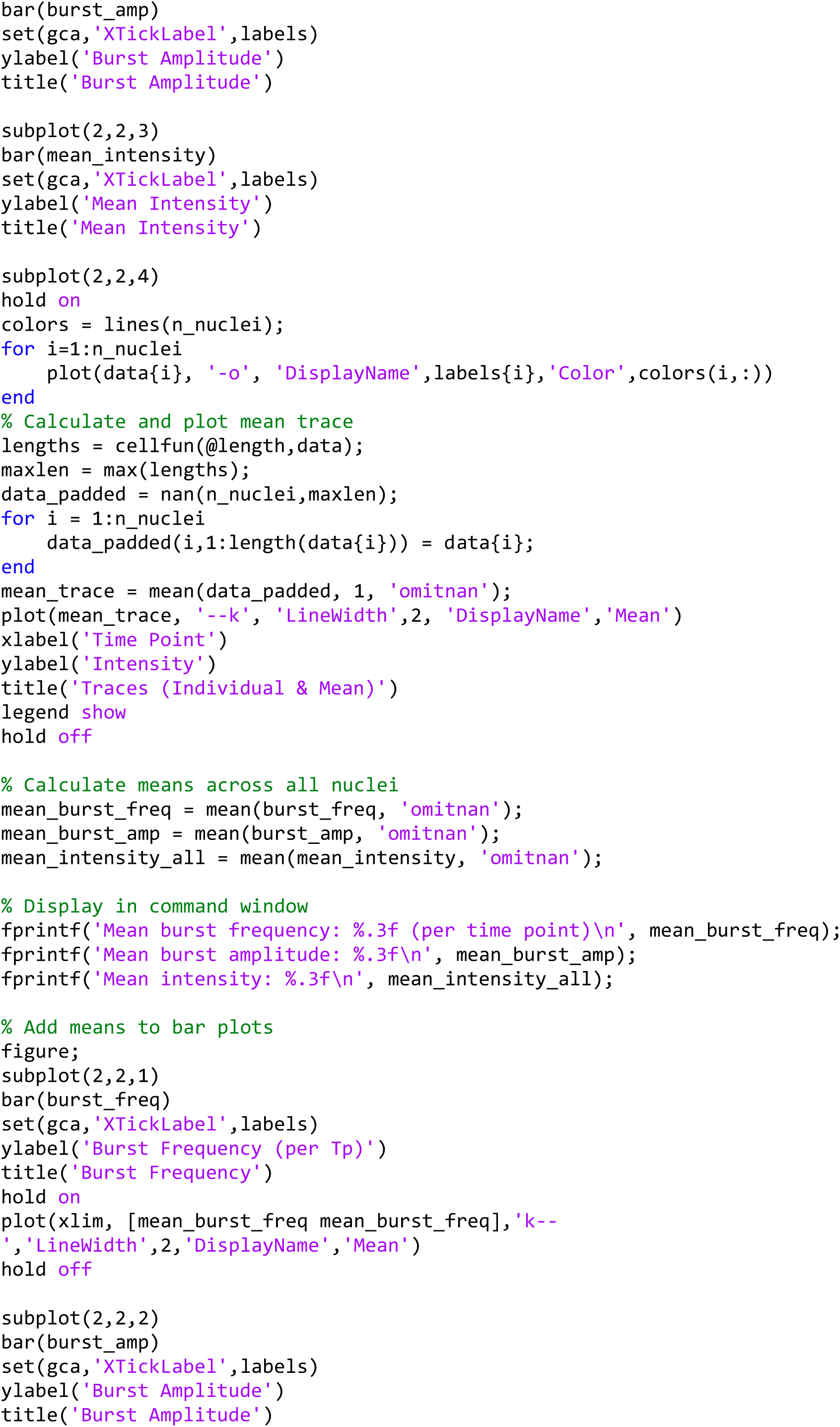
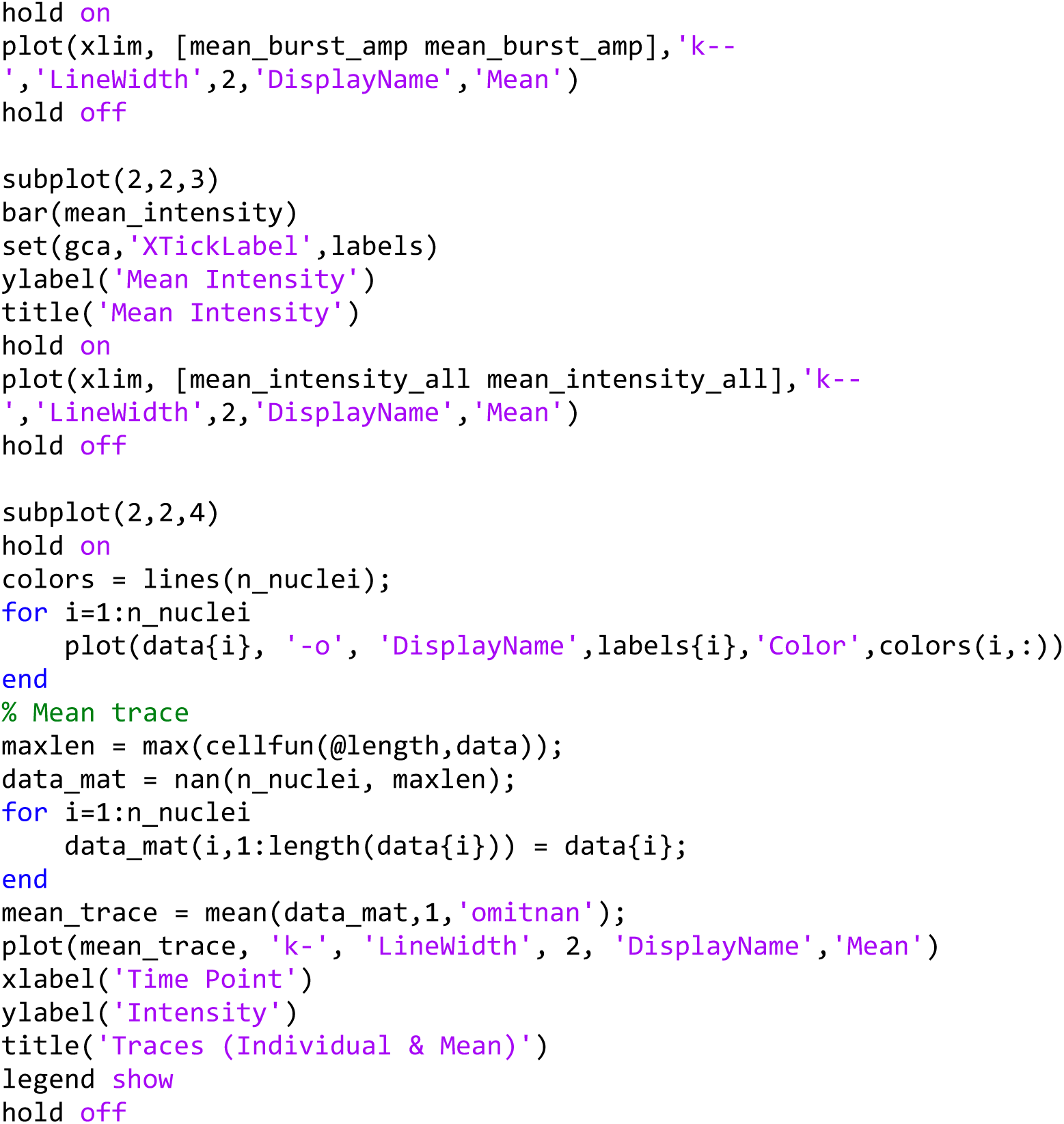

